# Personalized Genome-Scale Metabolic Models Identify Targets of Redox Metabolism in Radiation-Resistant Tumors

**DOI:** 10.1101/2020.04.07.029694

**Authors:** Joshua E. Lewis, Tom E. Forshaw, David A. Boothman, Cristina M. Furdui, Melissa L. Kemp

## Abstract

Redox cofactor production is integral towards antioxidant generation, clearance of reactive oxygen species, and overall tumor response to ionizing radiation treatment. To identify systems-level alterations in redox metabolism which confer resistance to radiation therapy, we developed a bioinformatics pipeline for integrating multi-omics data into personalized genome-scale flux balance analysis models of 716 radiation-sensitive and 199 radiation-resistant tumors. These models collectively predicted that radiation-resistant tumors reroute metabolic flux to increase mitochondrial NADPH stores and ROS scavenging. Simulated genome-wide knockout screens agreed with experimental siRNA gene knockdowns in matched radiation-sensitive and –resistant cancer cell lines, revealing gene targets involved in mitochondrial NADPH production, central carbon metabolism, and folate metabolism that allow for selective inhibition of glutathione production and H_2_O_2_ clearance in radiation-resistant cancers. This systems approach represents a significant advancement in developing quantitative genome-scale models of redox metabolism and identifying personalized metabolic targets for improving radiation sensitivity in individual cancer patients.

## Introduction

Radiation therapy remains a cornerstone of cancer treatment, with more than half of all cancer patients receiving radiation as part of their treatment regimen (Delaney et al., 2005; Miller et al., 2016). Nonetheless, tumor resistance to radiation therapy constitutes a significant obstacle to long-term cancer patient survival. More than one-fifth of patients in The Cancer Genome Atlas (TCGA) database continued to show stable or progressive disease following radiation treatment, and almost all cancer types had some proportion of radiation-resistant patients (Weinstein et al., 2013). To investigate the underlying pathophysiological mechanisms of radiation resistance and discover targets for improving sensitivity to radiation therapy, correlative studies using single omics modalities such as genomics or transcriptomics have been previously performed (Chen et al., 2015; Lee et al., 2010; Manem and Dhawan, 2019; Skvortsov et al., 2014; Skvortsova et al., 2008; Smith et al., 2009). However, these previous findings fall short by failing to integrate multiple biological data types, analyze differential expression in the context of genome-scale biochemical and regulatory networks, or provide mechanistic insights into how prospective biomarkers impact tumor function and can be exploited to improve radiation sensitivity.

Redox metabolism relies on the oxidation and reduction of electron-carrying molecules such as NADPH, NADH, and glutathione (GSH), which are used as cellular antioxidants and electron donors for metabolic reactions (Lewis et al., 2019; Xiao et al., 2018). These metabolites represent the reduced forms of redox couples with their associated oxidized forms NADP^+^, NAD^+^, and glutathione disulfide (GSSG), respectively; the ratio of reduced to oxidized forms of these redox couples provides an indication of the intracellular redox potential and oxidative state of the cell (Mallikarjun et al., 2012). Ionizing radiation therapy results in the generation of reactive oxygen species (ROS) such as superoxide (O_2_^-^) and hydrogen peroxide (H_2_O_2_), which oxidize the cellular environment and damage cellular structures including DNA (Brady et al., 2013; Cadet and Wagner, 2013; Reisz et al., 2014; Tominaga et al., 2004). Redox cofactors such as NADPH and GSH can be utilized by H_2_O_2_-scavenging enzymes to lower cellular levels of ROS (Forshaw et al., 2019; Harris et al., 2015). Additionally, these cofactors can directly promote DNA damage repair following oxidative damage, either by reduction of nitrogenous bases or utilization of NAD(P)H for nucleotide synthesis (Alvarez-Idaboy and Galano, 2012; Chatterjee, 2013; Franklin et al., 2016; Turgeon et al., 2018). Since redox metabolism is critical to the response of tumors to ionizing radiation, identifying targets for inhibiting production of these redox cofactors may provide a valuable strategy for sensitizing tumors to radiation therapy (Lewis et al., 2019).

Because redox cofactors are utilized in thousands of reactions throughout the human metabolic network, computational methods are needed to investigate systems-level redox metabolism and its interconnections with other cellular metabolic pathways (Brunk et al., 2018). Flux balance analysis (FBA) is a computational approach for predicting steady-state metabolic fluxes at a genome scale for cells or tissues of interest (Orth et al., 2010). By combining the stoichiometric representation of the human metabolic network, constraints on the fluxes through metabolic reactions, and an objective function to maximize a particular metabolic phenotype, predictions of maximum reaction fluxes or metabolite production rates under physiological constraints are generated (Bordel, 2018; Oberhardt et al., 2010; Supandi and Van Beek, 2018). Recently, FBA models personalized to individual cancer cell lines or tumors have been developed through the integration of transcriptomic data; however, there remain significant methodological shortcomings in the construction of these models that have hindered their ability to yield accurate and quantitative metabolic predictions (Blazier and Papin, 2012). Commonly-used FBA algorithms such as GIMME, iMAT, MADE, and CORDA utilize arbitrary gene expression thresholds to constrain metabolic activity, necessitate the comparison of multiple transcriptomic datasets, or completely remove reactions from the metabolic network with low associated gene expression (Becker and Palsson, 2008; Jensen and Papin, 2011; Schultz and Qutub, 2016; Shlomi et al., 2008). Other methods such as E-Flux constrain maximum reaction fluxes in proportion to the associated enzyme’s gene expression, but arbitrary reference values and proportionality functions are still used to set flux constraints instead of directly estimating enzyme abundances (Colijn et al., 2009). Additionally, most FBA models fail to incorporate any kinetic or thermodynamic constraints, which greatly affect metabolic fluxes and the directionalities of individual reactions (Henry et al., 2007). These shortcomings in model construction have ultimately limited the clinical utility of FBA models for accurately predicting metabolic phenotypes and directly improving cancer diagnosis and treatment (Zhang and Hua, 2015).

We have resolved many of these limitations in FBA model development and utility by integrating transcriptomic, kinetic, and thermodynamic information into quantitative constraints on the maximum flux and directionality of metabolic reactions. In our previously-developed FBA models of radiation-sensitive and radiation-resistant head and neck squamous cell carcinoma (HNSCC) cell lines, we accurately identified oxidoreductase genes that differentially impacted response to treatment with the NADPH-dependent redox-cycling chemotherapeutic β-lapachone between radiation-sensitive and -resistant cells (Lewis et al., 2018). Additionally, our models suggested that radiation-resistant cancer cells re-route NADH-generating metabolic fluxes through NAD salvage and purine salvage pathways involving NAMPT, an enzyme whose activity has previously been associated with radiation resistance and poor survival in cancer patients (Gujar et al., 2016; Lewis et al., 2019). Here, we extend this approach by developing an automated bioinformatics pipeline for integrating multi-omics information from The Cancer Genome Atlas (TCGA) and publically-available repositories into personalized genome-scale FBA models of 915 patient tumors across multiple cancer types. These personalized FBA models are used to investigate differences in redox cofactor production between radiation-sensitive and -resistant tumors, discover gene targets which differentially impact redox metabolism in radiation-resistant tumors, and identify personalized therapeutic strategies for individual radiation-resistant patients.

## Results

### An automated bioinformatics pipeline integrates multi-omics data into personalized FBA models of TCGA patient tumors

The framework for building FBA models of tumor metabolism was initiated with the community-curated Recon3D human metabolic reconstruction (8,401 metabolites, 13,547 reactions, and 3,268 genes; **Figure 1A**) (Brunk et al., 2018). The stoichiometric representation of this reconstruction is combined with minimum and maximum constraints on reaction fluxes to obtain a solution space for steady-state fluxes throughout the metabolic network. An objective function is typically utilized to narrow the possible FBA solution space to physiologically-optimal metabolic fluxes (Garcia Sanchez and Torres Saez, 2014). We hypothesized that tumors are under selective pressure to maximize production of reduced redox cofactors including NADPH, NADH, and GSH to decrease ROS-mediated damage induced by ionizing radiation; thus, we used an objective function of maximizing redox cofactor reduction. FBA is used to compare maximum cofactor reduction between radiation-sensitive and -resistant tumors, while flux variance analysis (FVA) is used to predict fluxes through individual metabolic reactions involved in cofactor reduction (**Figure 1B**).

**Figure 1.**
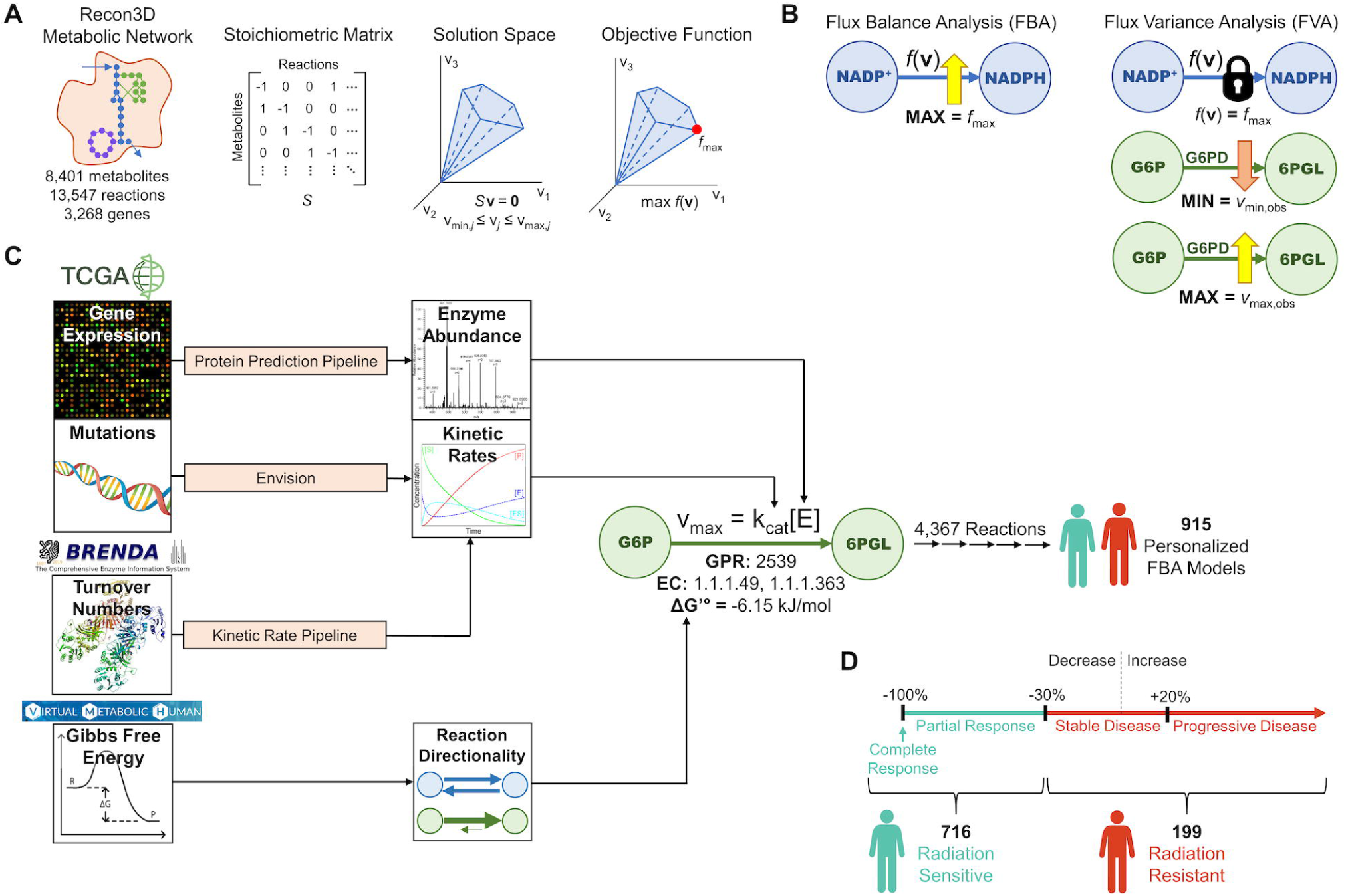
An automated bioinformatics pipeline integrates multi-omics data into personalized FBA models of TCGA patient tumors. (**A**) Implementation of flux balance analysis (FBA), including utilization of a stoichiometric representation of the Recon3D human metabolic network reconstruction, application of reaction constraints to obtain a solution space of flux values, and maximization of an objective function within this valid solution space. (**B**) Flux balance analysis (FBA) calculates the objective value, i.e., the maximum value of the objective function. Flux variance analysis (FVA) calculates the minimum and maximum possible fluxes through each metabolic reaction while maintaining the objective function at its maximum value. (**C**) Pipeline for integrating multi-omics data from The Cancer Genome Atlas (TCGA) and publically-available repositories into personalized FBA models of TCGA patient tumors. (**D**) Classification of TCGA patient tumors into radiation-sensitive and -resistant classes based on observed decrease/increase in tumor size following radiation therapy. See also **Figures S1-S2**.

To generate personalized FBA models of radiation-sensitive and -resistant TCGA patient tumors, Michaelis-Menten V_max_ constraints were set on all 4,367 Recon3D reactions that contain both a gene-protein-reaction (GPR) rule and enzyme commission (EC) number (**Figure 1C**). This constraint sets the maximum flux for each reaction (V_max_, units of mmol gDW^-1^ hr^-1^) equal to the kinetic rate constant of the enzyme catalyzing the reaction (k_cat_, units of hr^-1^), multiplied by the estimated protein abundance of the enzyme ([E], units of mmol gDW^-1^). A custom protein prediction pipeline was developed to convert RNA-seq transcriptomic data from individual TCGA samples into estimated enzyme abundances ([E]) (**Figure S1A**). This pipeline takes advantage of models relating the mRNA and protein abundances for individual genes/proteins using experimentally-measured transcription, translation, and degradation rates (Schwanhausser et al., 2011). Predicted enzyme abundances from this pipeline had improved correlation with experimental protein expression values from both NCI-60 and TCGA datasets compared to original gene expression values (**Figure S1B-I**). Additionally, a custom kinetic rate pipeline was developed to extract physiologically-accurate turnover numbers for metabolic enzymes (k_cat_) by matching experimentally-measured values from the BRENDA database with the correct enzyme, substrate, organism, and environmental conditions as those in the Recon3D network (**Figure S2**) (Schomburg et al., 2004). Envision database scores were applied to predict the effect of mutations in individual TCGA samples on the catalytic rate of corresponding metabolic enzymes (Gray et al., 2018). Finally, standard transformed changes in Gibbs free energy (ΔG’°) from the Virtual Metabolic Human (VMH) database were used to set thermodynamic constraints, such that only reactions with a negative ΔG^’°^ can carry non-zero net fluxes (Noor et al., 2013). These proteomic, kinetic, and thermodynamic constraints yield quantitative, patient-specific, and physiologically-accurate predictions of metabolic fluxes on a genome scale.

RECIST classification of TCGA samples provided an evaluation metric of radiation sensitivity based on changes in tumor size in response to radiation therapy (**Figure 1D**). Patients with a complete or partial response to radiation (greater than 30% decrease in tumor size) were classified as radiation-sensitive, and patients with stable or progressive disease (either less than 30% decrease in tumor size, or increase in tumor size) were classified as radiation-resistant. Using this classification, 716 personalized FBA models of radiation-sensitive tumors and 199 personalized models of radiation-resistant tumors were generated.

### Radiation-resistant tumors display compartmental differences in redox metabolic fluxes compared to radiation-sensitive tumors

Personalized FBA models were first used to compare differences in redox cofactor production between radiation-sensitive and -resistant tumors. Radiation-resistant tumor models showed significantly elevated production of reduced cofactors NADPH, NADH, and GSH (**Figure 2A**). To validate model predictions, we performed experimental measurements in matched pairs of radiation-sensitive and -resistant cell lines across three different cancer types (**Figure 2B, Table S1**). Two of the three radiation-resistant cell lines had significantly more reduced glutathione half-cell potentials (E_hc_ GSH/GSSG) than their matched radiation-sensitive cell line, indicating greater conversion of oxidized GSSG to reduced GSH as predicted in TCGA models (**Figure 2C**). Although BRCA cell lines showed the opposite trend, the observed E_hc_ difference was much smaller than in the other two cancer types and could be attributed to compensation for *NQO1* depletion by increased expression of other antioxidant enzymes involved in glutathione reduction or which bypass the need for GSH (e.g. peroxiredoxin or thioredoxin systems) (Cao et al., 2014). While predicted cytosolic production of NADPH did not differ between tumor classes, increased mitochondrial NADPH production in radiation-resistant tumors accounted for observed differences in total cellular NADPH production (**Figure 2D**). In agreement with this finding, increased deoxynucleotide production (which relies on mitochondrial NAD(P)H) was seen in radiation-resistant tumor models, while no significant differences in production of fatty acid precursors including palmitate (which relies on cytosolic NAD(P)H) were observed (**Figure S3A-B**) (Jones, 1980; Lewis et al., 2014; Schnell et al., 2004; Turgeon et al., 2018).

**Figure 2.**
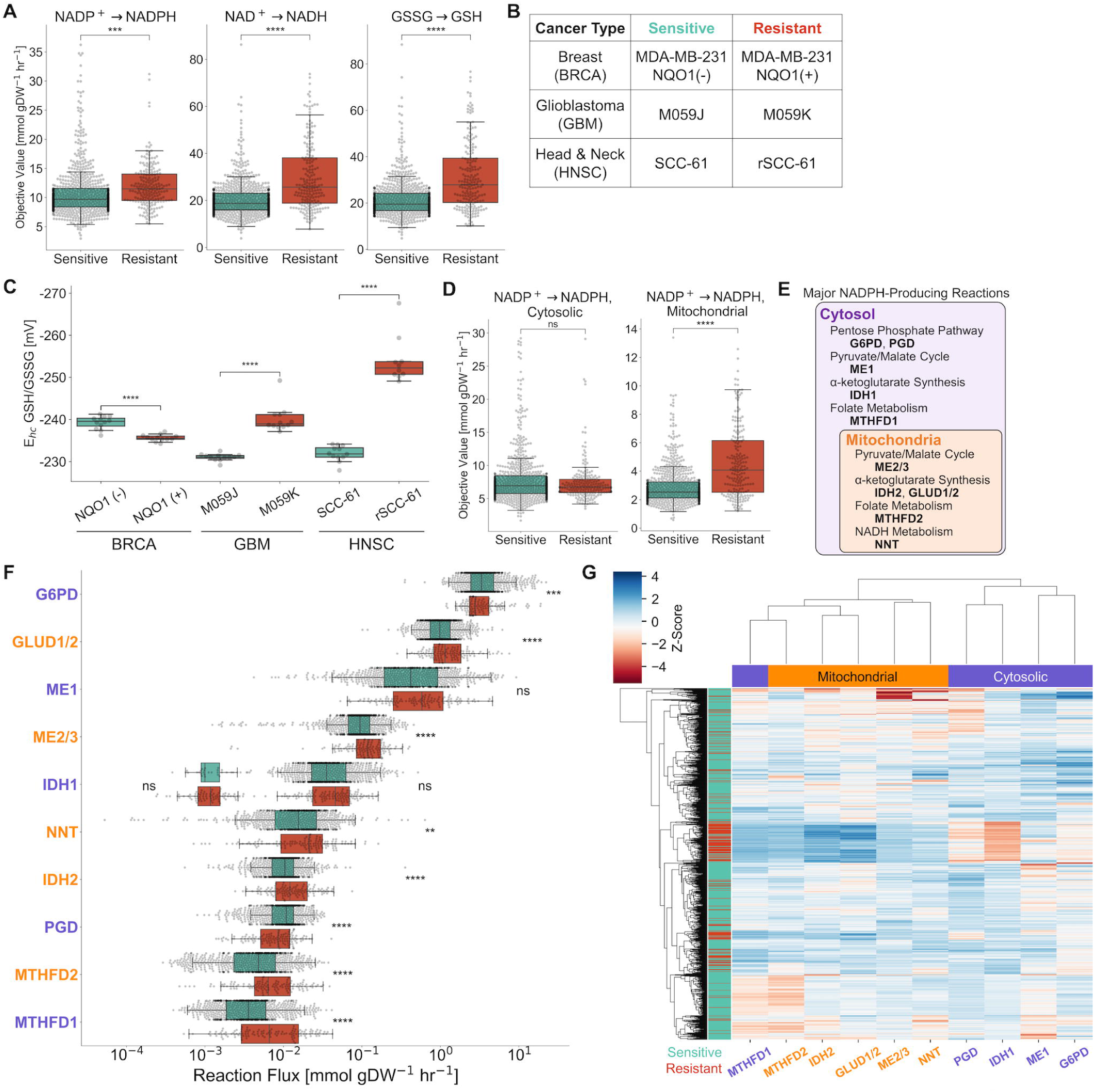
Radiation-resistant tumors display compartmental differences in redox metabolic fluxes compared to radiation-sensitive tumors. (**A**) Comparison of FBA-predicted production of reduced NADPH, NADH, and GSH between radiation-sensitive and -resistant TCGA tumors. (**B**) Matched pairs of radiation-sensitive and -resistant cell lines across multiple cancer types for experimental validation. (**C**) Experimentally-measured glutathione half-cell potential in cancer cell lines. (**D**) Comparison of FBA-predicted cytosolic and mitochondrial production of reduced NADPH. (**E**) Major NADPH-producing reactions with their associated cellular compartments and metabolic pathways. (**F**) FVA-predicted fluxes through major NADPH-producing reactions. IDH1 fluxes were separated between tumors with IDH1 R132 mutations (left) and wild-type IDH1 tumors (right). Reaction names are colored based on cellular compartment. (**G**) Hierarchical clustering of FVA-predicted fluxes based on TCGA patient tumor (rows) and NADPH-producing reaction (columns). Values are the Z-score of reaction fluxes across all tumors for each individual reaction. ns: not significant, *: p ≤ 0.05, **: p ≤ 0.01, ***: p ≤ 0.001, ****: p ≤ 0.0001. See also **Figure S3, Table S1**.

Flux variance analysis (FVA) was used to compare fluxes through major NADPH-generating metabolic reactions between radiation-sensitive and -resistant tumors (**Figure 2E-F**). Predicted fluxes through major cytosolic reactions including G6PD and PGD were greater in radiation-sensitive tumors, in agreement with our previously identified flux distributions in radiation-sensitive HNSCC cell lines as well as experimental findings (Lewis et al., 2018; Mims et al., 2015). On the other hand, predicted fluxes through major mitochondrial and folate-dependent reactions including GLUD1/2, ME2/3, NNT, IDH2, MTHFD2, and MTHFD1 were greater in radiation-resistant tumors, accounting for net increased NADPH production in these tumors. Hierarchical clustering of NADPH-producing fluxes yielded pronounced separation of cytosolic and mitochondrial reactions except for the folate-dependent reactions MTHFD1 and MTHFD2, both of which displayed increased fluxes in radiation-resistant tumor models (**Figure 2G**). Clustering of samples based on radiation response was found to be optimal compared to other clinical factors including cancer type as measured by the silhouette coefficient (**Figure S3C**). Collectively, these results suggest that differences in mitochondrial- and folate-dependent NADPH metabolism can discriminate between radiation-sensitive and -resistant tumors.

### Simulated genome-wide knockout screen identifies targets of redox metabolism in radiation-resistant cancers

FBA models enable the assessment of changes in metabolic fluxes in response to knockout of a particular metabolic enzyme-encoding gene, providing insight into the gene’s role within genome-scale metabolism (**Figure 3A**). A simulated genome-wide knockout screen was performed to predict the effect of knocking out each individual gene in Recon3D on total cellular NADPH production across all TCGA models (**Figure 3B**). Most gene knockouts did not significantly decrease total NADPH production, corroborating our previous findings that metabolic networks redirect flux through alternate, compensatory pathways following perturbation to optimize NADPH production (Lewis et al., 2018). Among the knockouts with largest predicted effects were genes directly involved in NADPH generation (*G6PD, GLUD1, ME1*), as well as those involved in glycolysis (*ALDOA, ENO1, GAPDH, GPI, HK1, PGAM1, PKM*), folate metabolism (*DHFR*), and amino acid metabolism (*PYCR2*).

**Figure 3.**
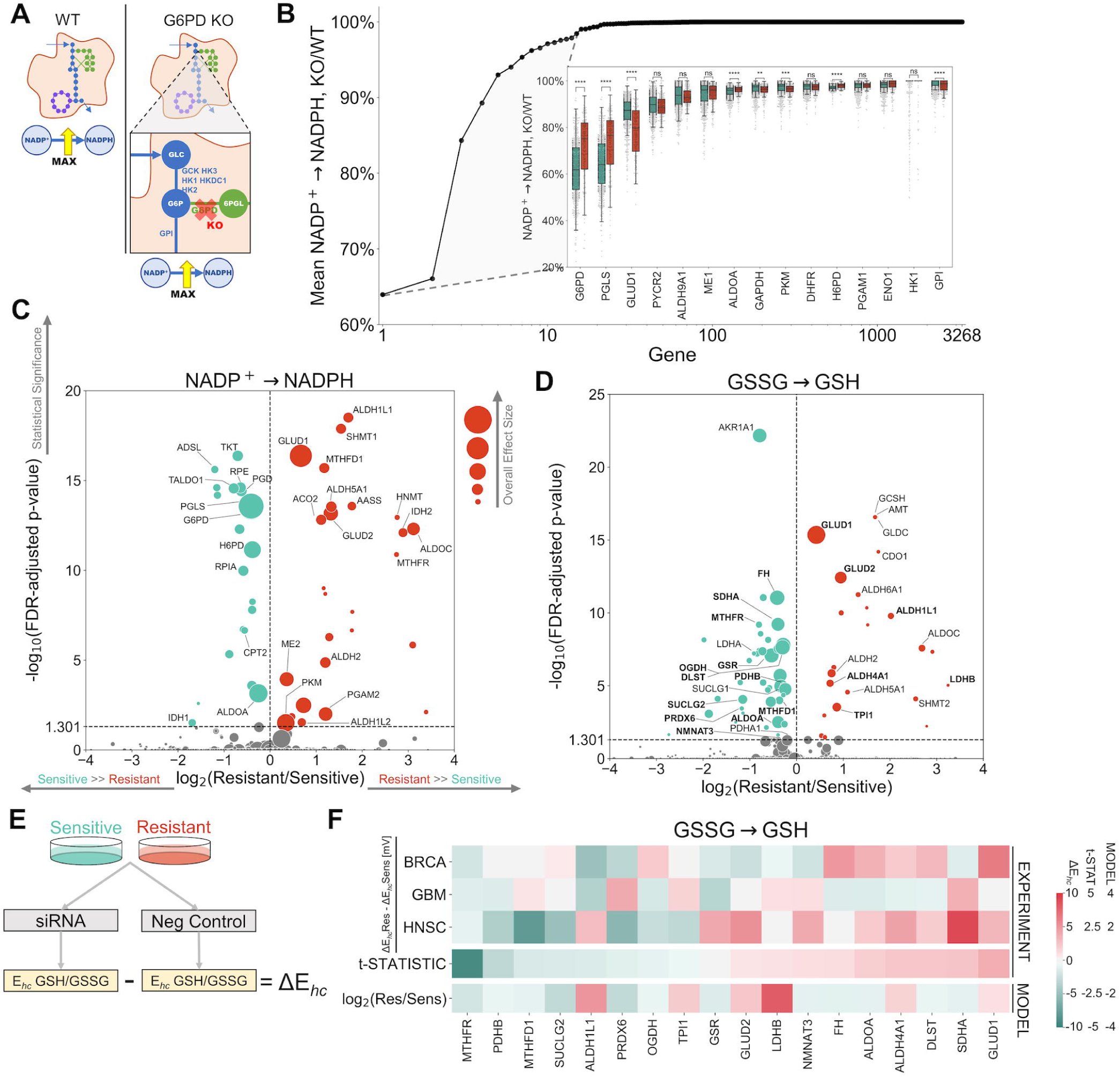
Simulated genome-wide knockout screen identifies targets of redox metabolism in radiation-resistant cancers. (**A**) Schematic comparing maximum total NADPH production between WT (left) and *G6PD*-knockout (right) models for individual patient tumors. Note that for reactions with more than one associated gene (e.g. GLUD1/2 reaction), only one gene is knocked out at a time. (**B**) Effect of simulated knockout of each individual gene in Recon3D on total NADPH production in TCGA tumors. Values are the ratio of total NADPH production after versus before knockout. Genes are rank ordered based on increasing mean KO/WT ratio (decreasing gene knockout effect) across all tumor models. Outset: KO/WT ratios are averaged across all tumor models. Inset: For the top 15 genes, KO/WT ratios from individual patient tumor models are shown, along with the comparison between radiation-sensitive and - resistant cohorts. (**C**) Volcano plot comparing the effect of each simulated gene knockout (individual dot) on total NADPH production between radiation-sensitive and - resistant tumors. X-axis: log_2_(Resistant/Sensitive), where “Resistant” equals the mean (WT-KO)/WT ratio in radiation-resistant tumors, and “Sensitive” equals the mean (WT-KO)/WT ratio in radiation-sensitive tumors; values < 0 (green dot on the left of the dotted line) signify knockouts with greater effects on total NADPH production in radiation-sensitive tumors, whereas values > 0 (red dot on the right of the dotted line) signify knockouts with greater effects on total NADPH production in radiation-resistant tumors. Y-axis: statistical significance (false discovery rate-adjusted p-values based on the Benjamini-Hochberg procedure) comparing knockout effects between radiation-sensitive and -resistant tumors; values above the dotted line (FDR-adjusted p-value ≤ 0.05) are statistically-significant. The size of each dot is proportional to the overall effect size (mean (WT-KO)/WT ratio across all tumor models regardless of radiation sensitivity). (**D**) Volcano plot comparing the effect of each simulated gene knockout on total reduced GSH production between radiation-sensitive and -resistant tumors. Genes tested by experimental siRNA knockdown studies are bolded. (**E**) Schematic demonstrating the measurement of ΔE_hc_ GSH/GSSG (difference in glutathione half-cell potential between siRNA knockdown and negative control) in radiation-sensitive and - resistant cancer cell lines. (**F**) Comparison of model-predicted and experimentally-measured effects of gene knockdown on reduced GSH production. Top 3 rows: ΔE_hc_Res - ΔE_hc_Sens in siRNA knockdowns across all three cell line pairs. ΔE_hc_Res - ΔE_hc_Sens > 0 for gene knockdowns causing greater oxidation in the radiation-resistant cell line, corresponding to a model-predicted log_2_(Resistant/Sensitive) > 0. Middle row: t-statistic from 1-sample t-test comparing the three experimentally-measured values of ΔE_hc_Res - ΔE_hc_Sens to the null hypothesis population mean of zero (equal effect in radiation-sensitive and -resistant cell lines). Bottom row: model-predicted log fold change in gene knockout effect on reduced GSH production between radiation-resistant and -sensitive TCGA tumor models. ns: not significant, *: p ≤ 0.05, **: p ≤ 0.01, ***: p ≤ 0.001, ****: p ≤ 0.0001. See also **Figure S4A**.

A statistical comparison of gene knockout effect on NADPH production between radiation-sensitive and -resistant tumors was performed to identify redox metabolic targets selective for radiation-resistant tumors (**Figure 3C**). 26 gene knockouts caused a significantly greater decrease in NADPH production among radiation-resistant tumors, including those involved in mitochondrial and folate-dependent NADPH metabolism (*GLUD1/2, IDH2, ME2, MTHFD1, MTHFR*) as well as many aldehyde dehydrogenase genes (*ALDH1L1, ALDH1L2, ALDH5A1*); many of these targets are consistent with those previously implicated in the response of radiation-resistant HNSCC to NADPH-depleting chemotherapies (Lewis et al., 2018). 24 gene knockouts caused a significantly greater decrease in NADPH production among radiation-sensitive tumors, including those of the pentose phosphate pathway (*G6PD, PGD, PGLS, RPE, RPIA, TALDO1, TKT*). While many targets with differential effects on NADPH production were also found to differentially impact GSH production between radiation-sensitive and -resistant tumor models, some notable differences were observed (**Figure 3D, Figure S4A**). Whereas pentose phosphate pathway genes were not predicted to significantly affect GSH production, those involved in the TCA cycle (*DLST, FH, OGDH, SDHA, SUCLG2*), oxidation-reduction reactions (*AKR1A1, GSR, PRDX6*), and glycine metabolism (*AMT, GCSH, GLDC*) were identified; glycine metabolism has been previously shown to impact GSH/GSSG ratios and ROS levels in cancer (Zhuang et al., 2018). Additionally, folate metabolism genes *MTHFD1* and *MTHFR* had larger impacts on NADPH production in radiation-resistant tumor models but impacted GSH production to a greater extent in radiation-sensitive models, possibly attributing to folate and NADPH’s role in other metabolic pathways including nucleotide synthesis (Fan et al., 2014).

To validate model-predicted targets of glutathione metabolism, the change in glutathione half-cell potential (ΔE_hc_ GSH/GSSG) between siRNA knockdown cells and negative control siRNA-transfected cells was measured in both radiation-sensitive and - resistant cell lines for 18 different gene targets (**Figure 3E**). 4 out of 6 model-predicted radiation-resistant targets (*ALDH4A1, GLUD1, GLUD2, LDHB*) had an overall more oxidizing effect in radiation-resistant cell lines, with *GLUD1* having the largest differential oxidizing effect across all siRNA’s tested (**Figure 3F**). Additionally, the 4 radiation-sensitive targets with the largest predicted differential effect size (*GSR, PRDX6, MTHFR*, and *SUCLG2*) all had an overall more oxidizing effect in radiation-sensitive cell lines. These results suggest that FBA models of TCGA tumors can be used to predict gene targets which differentially impact redox cofactor production associated with radiation sensitivity.

### Disparities in redox metabolism and H_2_O_2_-scavenging systems between radiation-sensitive and -resistant tumors determine differential ROS response

By utilizing an FBA objective function maximizing H_2_O_2_ clearance, radiation-resistant tumor models predicted increased H_2_O_2_-scavenging potential compared to radiation-sensitive tumors (**Figure 4A**). To validate model predictions of H_2_O_2_ response, matched radiation-sensitive and -resistant cell line pairs were treated for 2 hours with 10 mU/mL of the H_2_O_2_-generating enzyme glucose oxidase (Adimora et al., 2010; Bankar et al., 2009; Daniela et al., 2015). All cell lines showed decreased viability with glucose oxidase treatment, but 2 out of the 3 radiation-resistant cell lines showed a significantly lesser decrease in relative cell viability compared to their matched radiation-sensitive cell lines (**Figure 4B**). These findings were consistent with experimentally-measured differences in glutathione half-cell potential (**Figure 2C**). FVA was used to compare fluxes through major H_2_O_2_-clearing reactions between radiation-sensitive and -resistant tumors, accounting for enzyme isoform differences between cellular compartments (**Figure 4C**). Mitochondrial H_2_O_2_-clearing fluxes through catalase (CAT), glutathione peroxidase (GPx) and glutaredoxin (Grx) were significantly greater in radiation-resistant tumor models, in agreement with previous model predictions of increased mitochondrial NADPH production.

**Figure 4.**
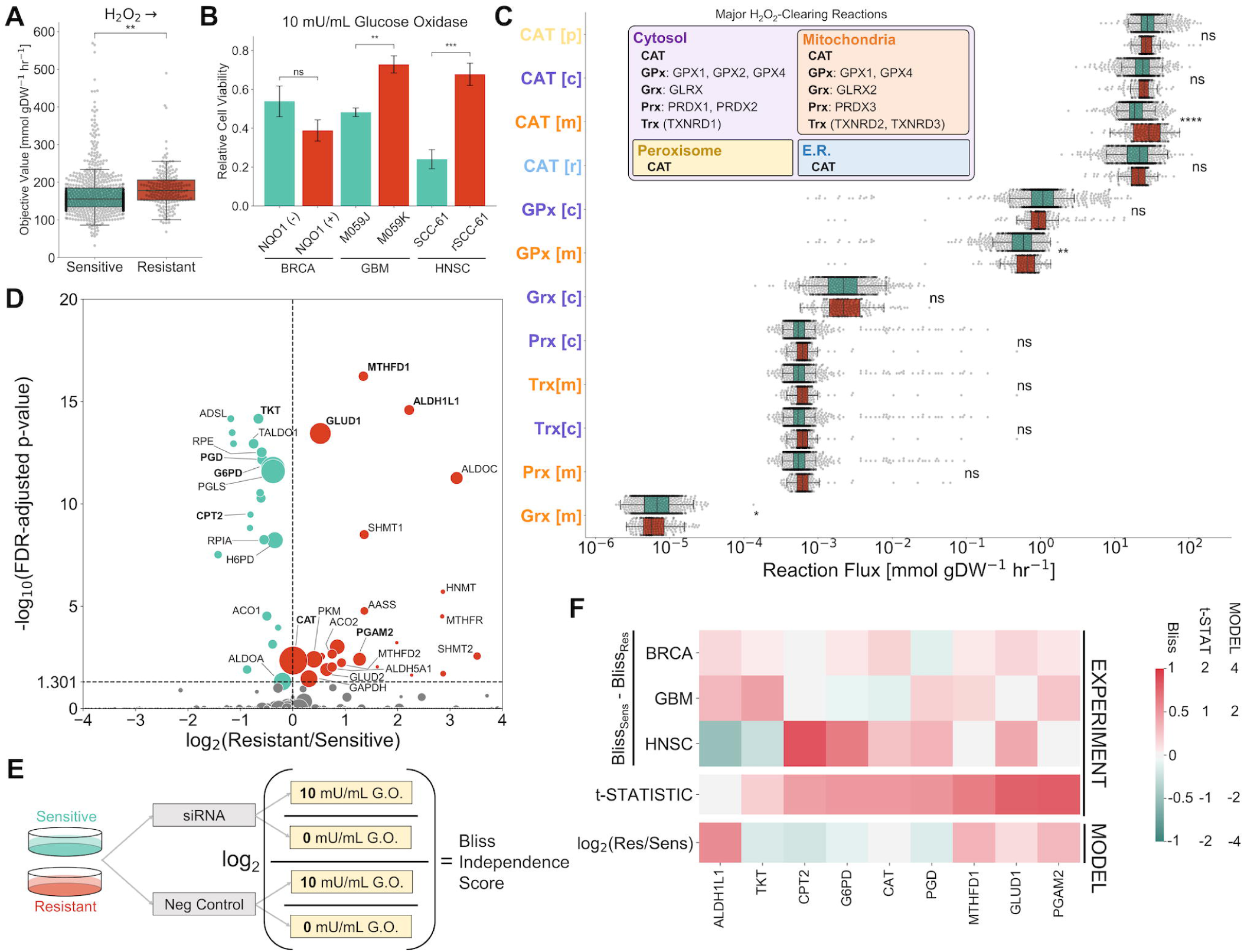
Disparities in redox metabolism and H_2_O_2_-scavenging systems between radiation-sensitive and -resistant tumors determine differential ROS response. (**A**) Comparison of FBA-predicted total clearance of H_2_O_2_ between radiation-sensitive and -resistant TCGA tumors. (**B**) Experimentally-measured response of radiation-sensitive and -resistant cancer cell lines to 2 hr treatment of 10 mU/mL glucose oxidase, calculated as the relative cell viability compared to 0 mU/mL glucose oxidase. Data are represented as mean ± 1 standard error. (**C**) FVA-predicted fluxes through major H_2_O_2_-clearing reactions. Inset: Major H_2_O_2_-clearing reactions with their associated cellular compartments and compartment-specific isoforms. (**D**) Volcano plot comparing the effect of each gene knockout on total H_2_O_2_ clearance between radiation-sensitive and - resistant tumors. Genes tested by experimental siRNA knockdown studies are bolded. **(E)** Schematic showing the measurement of Bliss independence scores in radiation-sensitive and -resistant cancer cell lines. (**F**) Comparison of model-predicted and experimentally-measured effects of gene knockdown on H_2_O_2_ clearance. Top 3 rows: Bliss_Sens_ - Bliss_Res_ in siRNA knockdowns across all three cell line pairs. Bliss_Sens_ - Bliss_Res_ > 0 for gene knockdowns causing a greater decrease in cell viability with glucose oxidase treatment in the radiation-resistant cell line, corresponding to a model-predicted log_2_(Resistant/Sensitive) > 0. Middle row: t-statistic from 1-sample t-test comparing the three experimentally-measured values of Bliss_Sens_ - Bliss_Res_ to the null hypothesis population mean of zero (equal effect in radiation-sensitive and -resistant cell lines). Bottom row: model-predicted log fold change in gene knockout effect on H_2_O_2_ clearance between radiation-resistant and -sensitive TCGA tumor models. ns: not significant, *: p ≤ 0.05, **: p ≤ 0.01, ***: p ≤ 0.001, ****: p ≤ 0.0001. See also **Figure S4B**.

A simulated genome-wide knockout screen was performed to identify gene targets with greater effects on H_2_O_2_ clearance in radiation-resistant tumors (**Figure 4D, Figure S4B**). While *CAT* was the only significant gene with direct H_2_O_2_-scavenging function, many gene targets of NADPH production were predicted to have significant differential effects on H_2_O_2_ clearance. Radiation-sensitive targets included members of the pentose phosphate pathway (*G6PD, PGD, PGLS, RPE, RPIA, TALDO1, TKT*) whereas radiation-resistant targets included genes involved in mitochondrial and folate-dependent NADPH production (*GLUD1/2, MTHFD1, MTHFD2, MTHFR*) and central carbon metabolism (*ACO2, ALDOC, GAPDH, PGAM2, PKM*). To validate these targets, Bliss independence scores indicating the effect of siRNA gene knockdown on glucose oxidase response were measured in radiation-sensitive and -resistant cell lines for 9 different gene targets (**Figure 4E**). 3 of the model-predicted radiation-resistant targets (*GLUD1, MTHFD1*, and *PGAM2*) had the largest differential response in radiation-resistant cell lines across all siRNA’s tested (**Figure 4F**). Furthermore, a larger differential response to ALDH1L1 knockdown was observed in 2 of the 3 radiation-resistant cell lines, as predicted by FBA models. TCGA models accurately predicted that *MTHFD1* knockdown would have a greater effect on GSH production in radiation-sensitive cancers but greater effect on H_2_O_2_ response in radiation-resistant cancers, suggesting that FBA models accurately capture other metabolic systems which impact ROS clearance besides glutathione-dependent pathways (Forshaw et al., 2019). Overall, the observed agreement between FBA model predictions and experimental validation demonstrates the ability to correctly identify targets of redox metabolism which differentially impact radiation-resistant cancers.

### Personalized metabolic flux profiles highlight heterogeneity in redox metabolism between radiation-resistant tumors

Although radiation-resistant tumor models displayed overall differences in redox metabolism compared to radiation-sensitive models, patient tumors may exemplify divergent metabolic flux profiles and thus differences in optimal therapeutic strategies for improving radiation sensitivity (Kim and DeBerardinis, 2019). To determine if patient clinical information could be used to distinguish radiation-resistant tumor models with differing metabolic phenotypes, we evaluated the correlation between clinical factors and predicted fluxes through major NADPH-generating reactions (**Figure 5A**). While cancer type, patient age, and tumor grade were highly correlated with predicted fluxes through most reactions, other clinical factors were associated with a few select reactions. Smoking history was highly associated with G6PD and PGD fluxes, in agreement with previous experimental studies showing that exposure to cigarette smoke causes upregulation of G6PD and shifts glucose metabolism towards the pentose phosphate pathway for increased NADPH production (Agarwal et al., 2014; Noronha-Dutra et al., 1993). Additionally, response to cisplatin treatment was highly associated with fluxes through mitochondrial reactions IDH2 and NNT; treatment with cisplatin has been previously reported to induce a mitochondrial-ROS response, which could lead to upregulation of mitochondrial NADPH-generating enzymes (Choi et al., 2015; Marullo et al., 2013).

**Figure 5.**
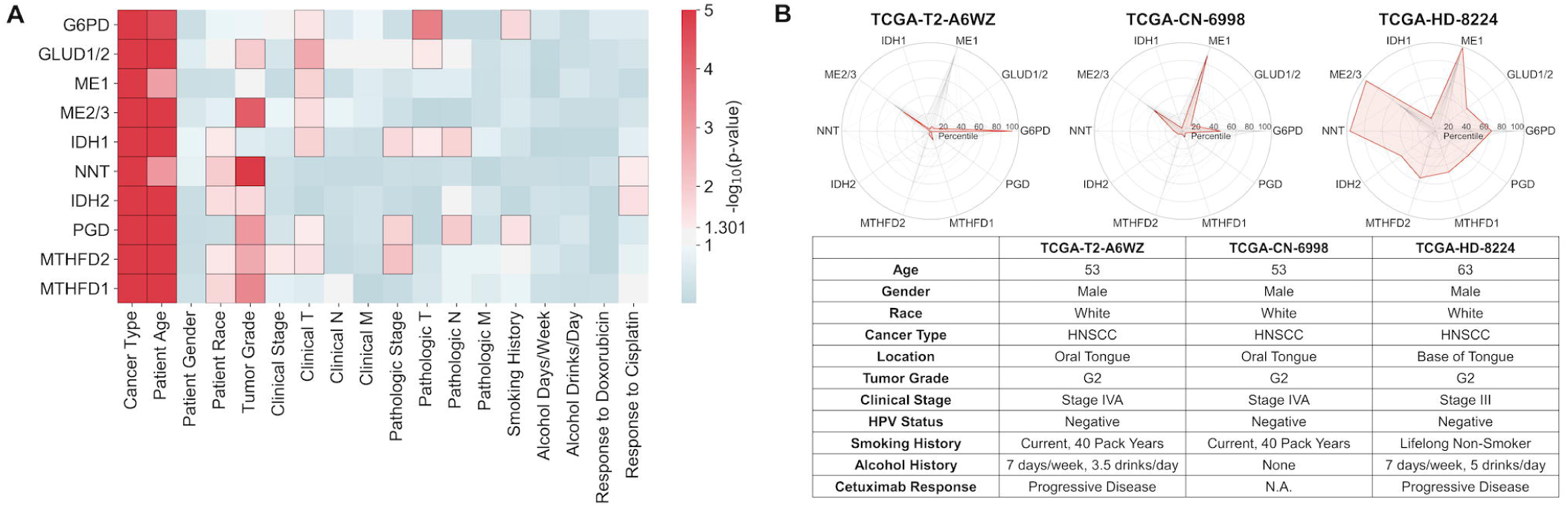
Personalized metabolic flux profiles highlight heterogeneity in redox metabolism between radiation-resistant tumors. (**A**) Correlation between patient clinical factors and predicted fluxes through major NADPH-producing reactions among radiation-resistant patients. Values are represented as the p-value of either the univariate regression (for numerical factors) or 1-way ANOVA (for categorical factors) between reaction fluxes and clinical factor values. Statistically significant (p ≤ 0.05) associations are represented with black borders. (**B**) Personalized NADPH-generating flux profiles of 3 radiation-resistant HNSCC patients with their associated clinical factors. Radar chart values are the percentiles of reaction fluxes across all radiation-resistant tumors for each individual reaction. Patient profiles (red, filled) are shown overlaid on top of the profiles of all other radiation-resistant HNSCC tumors (black, not filled).

To investigate whether patients with similar clinical features also displayed similarities in tumor redox metabolism, we evaluated personalized metabolic flux profiles of radiation-resistant patients with head and neck squamous cell carcinoma, a cancer type characterized by high genetic and metabolic heterogeneity (**Figure 5B**) (Chen et al., 2018). Predicted fluxes through NADPH-generating reactions differed substantially between patients with very similar clinical features; while some tumor models showed increased NADPH generation through single reactions (TCGA-T2-A6WZ: G6PD; TCGA-CN-6998: ME1), others showed increased fluxes through multiple disparate reactions (TCGA-HD-8224: ME1, ME2/3, NNT). These stark differences in redox metabolism among patients with similar clinical and demographic attributes demonstrate the utility of personalized genome-scale metabolic models for identifying therapeutic targets in individual patients.

## Discussion

Identifying redox metabolic targets which impact radiation sensitivity through modulation of antioxidant levels and oxidative DNA damage is an ongoing challenge (Spitz et al., 2004). Flux balance analysis (FBA) is a genome-scale metabolic modeling approach that has gained attraction for building mechanistic models of cancer metabolism and identifying chemotherapeutic targets (Folger et al., 2011; Nilsson and Nielsen, 2017; Yizhak et al., 2014). Because redox cofactors are involved in thousands of reactions throughout the human metabolic network, genome-scale FBA models are well-suited to study redox metabolism and their interconnections with other metabolic processes (Brunk et al., 2018). Nonetheless, few studies have used FBA models for studying cancer redox metabolism. A notable example is that of *Benfeitas et al*., who developed FBA models of hepatocellular carcinomas (HCC) to characterize heterogeneity in redox metabolism between HCC tumors at differing stages of progression and within different HCC tumor clusters (Benfeitas et al., 2019). Using these models, the authors discovered distinct differences in NADPH production and H_2_O_2_ clearance between HCC tumors with high *G6PD* expression and those with high *ALDH2* expression. However, their use of the MADE algorithm for converting transcriptomic data into upper flux bounds results in discrete reaction constraints which lack quantitative accuracy compared to using continuous enzyme abundance values. Furthermore, kinetic and thermodynamic effects on reaction constraints, which have significant impacts on metabolic fluxes, were not present in these models (Henry et al., 2007).

Our bioinformatics and metabolic modeling pipeline represents, to our knowledge, the first study to integrate multiple omics datasets into human genome-scale mechanistic models to compare redox metabolism between radiation-sensitive and -resistant tumors, as well as to identify therapeutic biomarkers for improving radiation therapy response. Methodological shortcomings of previous FBA studies are overcome by incorporating transcriptomic and mutational data from individual patient tumors, as well as genome-scale kinetic and thermodynamic parameter values, into quantitative constraints on metabolic fluxes, allowing for more accurate metabolic predictions (**Figure 1**). FBA models of radiation-resistant TCGA tumors showed increased fluxes through mitochondrial NADPH-producing reactions, allowing for elevated mitochondrial stores of reduced redox cofactors as well as increased fluxes through mitochondrial H_2_O_2_-clearing reactions (**Figures 2, 4**). Mitochondrial NADPH-producing enzymes identified from this study, including GLUD1, IDH2, ME2, and NNT, have been previously implicated in regulation of oxidative stress and tumor proliferation; additionally, mitochondrial-specific compartmentalization of signaling and energy metabolism has been previously identified to impact chemotherapy and radiation response (Ciccarese and Ciminale, 2017; Hsieh et al., 2015; Jin et al., 2015; Porporato et al., 2018; Stein et al., 2014; Yin et al., 2012). Simulated genome-wide knockout screens and validation with experimental siRNA knockdown experiments suggest that enzymes involved in mitochondrial (GLUD1/2) and folate-dependent (MTHFD1) NADPH production and central carbon metabolism (LDHB, PGAM2) may be viable targets for inhibiting GSH production and/or H_2_O_2_ clearance in radiation-resistant tumors (**Figures 3, 4**). On the other hand, G6PD, the most well-characterized NADPH-producing reaction, exemplified larger fluxes in radiation-sensitive tumor models and showed lower experimental effects on targeted H_2_O_2_ clearance in radiation-resistant cell lines compared to the aforementioned gene targets. Interestingly, the *G6PD* cluster identified by *Benfeitas et al*. in HCC is highly enriched in genes identified from this current study as radiation-sensitive targets (*ALDOA, G6PD, PGD, RPE, RPIA*), whereas the *ALDH2* cluster is highly enriched in genes identified as radiation-resistant targets (*ALDH2, ALDH5A1, ALDH6A1, CAT, MTHFD1, SHMT1*) (Benfeitas et al., 2019). Despite the fact that the authors were not analyzing redox metabolism in the context of radiation sensitivity, this correspondence between gene sets from separate analyses may suggest that there exist two fundamental tumor subtypes with distinct redox metabolic phenotypes and corresponding radiation sensitivities.

By developing FBA models of individual patient tumors, personalized metabolic flux profiles were generated to identify patient-specific redox biomarkers (**Figure 5**). These personalized metabolic predictions also demonstrate the significant amount of heterogeneity in redox metabolism between patient tumors. While the majority of initiatives for precision medicine for cancer treatment has focused on mutations and expression differences in signaling pathway proteins, there is currently a greater focus on exploiting metabolic differences between patient tumors (DeBerardinis and Chandel, 2016; Kanarek et al., 2018; Kanarek et al., 2020). For example, heterogeneity in glycolytic metabolism is being used to identify diagnostic biomarkers and treatment strategies for both pancreatic cancer and acute myeloid leukemia patients (Follia et al., 2019; Stuani et al., 2019). Personalized nutrition may also be a viable strategy for manipulating tumor redox metabolism in individual patients; most NADPH-generation pathways are supplied by glucose and glutamine intake, and folic acid levels may impact folate-dependent reactions (Choi and Park, 2018; Wallace et al., 2019). Continued development of genome-scale metabolic models of individual patient tumors will undoubtedly aid in the identification of metabolic targets or optimal dietary strategies for individual cancer patients.

Although our bioinformatics and modeling approach towards using multi-omics data for predicting metabolic phenotypes in individual patient tumors represents a significant methodological advancement over previous FBA models, additional improvements and integrations with other modeling strategies would further improve its accuracy and applicability. Currently, rates of metabolite transport through specific plasma membrane transporters are much less characterized than turnover rates of intracellular metabolic enzymes, limiting the implementation of quantitative transport constraints (Schomburg et al., 2004). Unless experimentally-measured metabolite uptake rates from samples of interest are obtained, constraints on uptake reactions are commonly set as binary (i.e. if a metabolite is present in the extracellular medium, intracellular uptake is unconstrained; otherwise, uptake is set to zero). A potential approach towards setting quantitative uptake constraints would be to relate membrane transporter expression with experimentally-measured extracellular metabolite concentrations (for example, from cell culture media or patient blood samples) to predict individualized metabolite uptake rates. In addition, regarding the application of FBA models towards studying redox metabolism, integration of important redox signaling and regulatory networks such as the Nrf2/Keap1/ARE pathway through methods such as integrated FBA (iFBA) could improve the accuracy of predicted reaction fluxes which utilize ARE-regulating genes including *G6PD, IDH1, ME1*, and *PGD* (Covert et al., 2008; Jaramillo and Zhang, 2013; Kansanen et al., 2013; Lee et al., 2008; Lin et al., 2016).

Despite the recent increase in experimental metabolomics studies towards identifying altered metabolic phenotypes in cancer, an integrated assessment of the 13,000+ human metabolic reactions and their collective impact on individualized treatment response is currently infeasible using solely experimental approaches (Kaushik and DeBerardinis, 2018). Instead, computational approaches which integrate multi-omics measurements and global reconstructions of human metabolism into predictive models provide tremendous utility in improving our understanding of pathophysiological processes and discovering personalized metabolic targets (Nilsson and Nielsen, 2017). Our analysis indicates that genome-scale metabolic models of individual patient tumors can identify important differences in redox metabolism between radiation-sensitive and -resistant tumors. Specifically, by comparing properties of 716 radiation-sensitive and 199 radiation-resistant personalized tumor models, we have elucidated multiple mechanisms of how tumors can upregulate metabolic flux through mitochondrial NADPH-generating and H_2_O_2_-clearing reactions to increase cellular antioxidant stores, decrease ROS levels, and resist the damaging effects of ionizing radiation treatment. These model-identified targets significantly impact antioxidant production and ROS response, thus serving as putative biomarkers for the *a priori* prediction of radiation sensitivity, as well as therapeutic strategies for sensitizing tumors to radiation therapy. Ultimately, the development of personalized metabolic models has the potential to facilitate the clinical management of cancer patients and improve long-term outcomes.

## Acknowledgements

The authors gratefully acknowledge support for this work from an NIH/NCI F30 CA224968 fellowship (PI: J.E.L.; Sponsor: M.L.K.) and an NIH/NCI U01 CA215848 grant (PIs: M.L.K., C.M.F., D.A.B.).

## Author Contributions

Conceptualization, J.E.L. and M.L.K.; Methodology, J.E.L.; Software, J.E.L.; Validation, J.E.L. and T.E.F.; Formal Analysis, J.E.L.; Investigation, J.E.L. and T.E.F.; Resources, M.L.K., C.M.F., and D.A.B.; Data Curation, J.E.L.; Writing – Original Draft, J.E.L.; Writing – Review & Editing, J.E.L., M.L.K., and C.M.F.; Visualization, J.E.L.; Supervision, M.L.K., C.M.F., and D.A.B.; Project Administration, J.E.L. and M.L.K.; Funding Acquisition, J.E.L. and M.L.K.

## Methods

### RESOURCE AVAILABILITY

### Lead Contact

Further information and requests for resources and reagents should be directed to and will be fulfilled by the Lead Contact, Melissa L. Kemp.

### Materials Availability

This study did not generate new unique reagents.

### Data and Code Availability

Code for the generation and simulation of personalized FBA models is available at https://github.com/kemplab/FBA-pipeline. Personalized models can be developed for any human sample (such as cell lines or patient tumors) with RNA-seq gene expression data, and mutation data if available. Jupyter notebooks are available for 1) processing of sample RNA-seq data to estimate enzyme abundance values, 2) processing of sample mutation data to estimate kinetic rate parameters, and 3) running FBA analysis with user-specified analysis type, model constraints, media constraints, objective function, and samples of interest. TCGA tumor models developed for this study are available as well.

The following datasets are available at https://github.com/kemplab/FBA-pipeline:

- *Dataset 1*. NADPH-generating fluxes for TCGA tumor models [mmol/gDW/hr] (Related to Figure 2)
- *Dataset 2*. Simulated effect of gene knockout on NADP^+^ → NADPH [KO/WT] (Related to Figure 3)
- *Dataset 3*. Simulated effect of gene knockout on GSSG → GSH [KO/WT] (Related to Figure 3)
- *Dataset 4*. Comparison of model-predicted and experimentally-validated GSSG → GSH gene targets (Related to Figure 3)
- *Dataset 5*. Simulated effect of gene knockout on H_2_O_2_ → [KO/WT] (Related to Figure 4)
- *Dataset 6*. Comparison of model-predicted and experimentally-validated H_2_O_2_ → gene targets (Related to Figure 4)

### EXPERIMENTAL MODEL AND SUBJECT DETAILS

#### Cell Lines

**Table S1** provides the matched radiation-sensitive and radiation-resistant human cancer cell lines used for experimental validation of model predictions. All cell lines were maintained in RPMI-1640 cell culture media (Thermo Fisher Scientific, Cat#11875) with 10% fetal bovine serum (Sigma-Aldrich, Cat#F4135) at 37°C and 5% CO_2_, and were free of *Mycoplasma*.

### METHOD DETAILS

#### Flux Balance Analysis (FBA)

A metabolic network can be represented by a stoichiometric matrix *S* of size *m* x *r*, where *m* and *r* are the number of metabolites and reactions in the network, respectively. Entry *S*_*ij*_ is equal to the stoichiometric coefficient of metabolite *i* in reaction *j* (*S*_*ij*_ < 0 for reactants, > 0 for products, and = 0 if metabolite *i* is not involved in reaction *j*). The relationship between reaction fluxes and metabolite concentrations in the network can be written as:

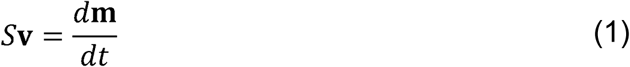

where **v** is a *r* x 1 vector of reaction fluxes, and **m** is a *m* x 1 vector of metabolite concentrations. In flux balance analysis, the steady state is assumed (metabolite concentrations do not change with time), changing Equation 1 to:

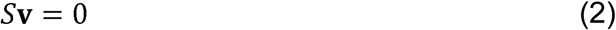

Each reaction flux *v*_*j*_ is also constrained by lower and upper bounds:

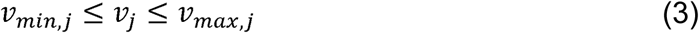

The solutions to Equations 2 and 3 that maximize a particular objective function *f*(**v**) are chosen. Thus, the flux balance analysis problem can be represented as a Linear Programming (LP) optimization problem:

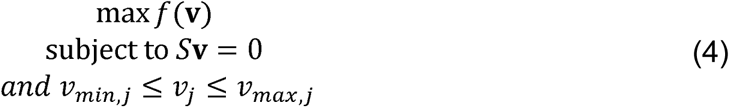

Solving Equation 4 provides the maximized value *f*_max_ of the objective function. Gurobi 8.0 optimization software was used to solve these LP problems.

#### Flux Variance Analysis (FVA)

Flux variance analysis allows for calculation of the minimum and maximum allowable fluxes through each metabolic reaction while still maintaining the maximum possible objective function value *f*_max_:

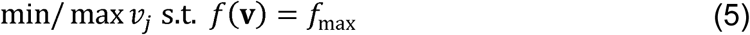

The average flux through each metabolic reaction was estimated by taking the mean of the minimum and maximum fluxes calculated from Equation 5.

#### Objective Functions

To maximize the production of a particular metabolite in the metabolic network, an artificial demand reaction can be added to the network, and the flux through this new objective function reaction can be maximized. This in turn maximizes the fluxes through other reactions throughout the metabolic network that produce the metabolite. FBA provides the objective value, i.e. the maximum amount of metabolite that can be produced, and FVA provides the fluxes through metabolic reactions which produce the metabolite. To maximize the reduction of a metabolite from its oxidized to reduced form, the objective function consists of an artificial demand reaction consisting of the oxidation of the metabolite - this in turn maximizes the fluxes through other metabolic reactions that reduce the metabolite.

For example, to maximize the reduction of NADP^+^ to NADPH in the cytosol, the objective function would be:

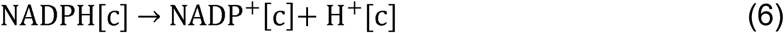

To maximize the production of a metabolite in all cellular compartments, separate objective functions for each compartment with the same weight were simultaneously maximized. **Table S2** lists the objective functions used in FBA and FVA. To ensure that biologically viable solutions were obtained, all models were checked to ensure that they were capable of producing physiological ATP levels typical of mammalian cells (1.0625 mmol gDW^-1^ hr^-1^) while maximizing the imposed objective functions (Kilburn et al., 1969).

#### Metabolic Network

Recon 3D version 3.01 was used as the core metabolic network (Brunk et al., 2018). While the “full” version (Recon3D_301.mat) produced erroneous energy-generating cycles (e.g. producing ATP without uptake of any metabolic precursors), the “model” version (Recon3DModel_301.mat) was found not to be the largest subset of reactions without erroneous energy-generating cycles (additional reactions from the “full” version could be added). To this end, following the procedure outlined in Fritzemeier et al., reactions from the “full” version were sequentially added back into the “model” version as long as they did not result in erroneous energy-generating cycles (Fritzemeier et al., 2017).

To address missing and inaccurate redox-based reaction information within Recon3D, the following changes were made:

1. The reaction “FTHFDH” was split into two separate cytosolic and mitochondrial reactions. The cytosolic reaction is catalyzed by ALDH1L1, and the mitochondrial reaction is catalyzed by ALDH1L2. *Originally:*
  ⍰ 1 10fthf[c] +1 h2o[c] + 1 nadp[c] → 1 co2[c] + 1 h[c] + 1 nadph[c] + 1 thf[c] GPR: 10840.1 or 160428.1 EC: 1.5.1.6
  *Updated*:
  ⍰ 1 10fthf[c] + 1 h2o[c] + 1 nadp[c] → 1 co2[c] + 1 h[c] + 1 nadph[c] + 1 thf[c] GPR: 10840.1 EC: 1.5.1.6
  ⍰ 1 10fthf[m] + 1 h2o[m] + 1 nadp[m] → 1 co2[m] + 1 h[m] + 1 nadph[m] + 1 thf[m] GPR: 160428.1 EC: 1.5.1.6
2. MTHFR was removed from the GPR associated with reaction “MTHFD”. MTHFR catalyzes the reaction converting 5,10-methylenetetrahydrofolate to 5-methyltetrahydrofolate, which is already included in Recon3D. *Originally*:
  ⍰ 1 mlthf[c] + 1 nadp[c] ← → 1 methf[c] + 1 nadph[c] GPR: 4522.1 or 4524.1 EC: 1.5.1.5
  *Updated*:
  ⍰ 1 mlthf[c] + 1 nadp[c] ← → 1 methf[c] + 1 nadph[c] GPR: 4522.1 EC: 1.5.1.5
3. Reactions catalyzed by isoforms of NADPH oxidase were added. *Originally*: None *Updated*:
  ⍰ 1 nadph[c] + 2 o2[c] → 2 o2s[e] + 1 nadp[c] + 1 h[c] GPR: 27035.1 or 1536.1 or 50508.1 or 79400.1 EC: 1.6.3.1
  ⍰ 1 nadph[n] + 2 o2[n] → 2 o2s[c] + 1 nadp[n] + 1 h[n] GPR: 50507.1 EC: 1.6.3.1
  ⍰ 1 nadph[r] + 2 o2[r] → 2 o2s[c] + 1 nadp[r] + 1 h[r] GPR: 50507.1 EC: 1.6.3.1
4. PRDX1 and PRDX2 were removed from the GPR associated with reaction “GTHP”. Glutathione peroxidase involves the reduction of hydrogen peroxide by glutathione, which does not involve peroxiredoxins. Peroxiredoxin reactions are added to Recon3D (see #7) *Originally*:
  ⍰ 1 h2o2[c] + 2 gthrd[c] → 2 h2o[c] + 1 gthox[c] GPR: 7001.3 or 5052.3 or 2877.1 or 2876.2 or 5052.2 or 2876.1 or 5052.1 or 7001.1 or 2879.1 or 7001.2 EC: 1.11.1.9
  *Updated*:
  ⍰ 1 h2o2[c] + 2 gthrd[c] → 2 h2o[c] + 1 gthox[c] GPR: 2877.1 or 2876.2 or 2876.1 or 2879.1 EC: 1.11.1.9
5. Similarly to #4, PRDX3 was removed from the GPR associated with reaction “GTHPm”. *Originally*:
  ⍰ 1 h2o2[m] + 2 gthrd[m] → 2 h2o[m] + 1 gthox[m] GPR: 10935.1 or 10935.2 or 2879.1 or 2876.1 EC: 1.11.1.9
  *Updated*:
  ⍰ 1 h2o2[m] + 2 gthrd[m] → 2 h2o[m] + 1 gthox[m] GPR: 2879.1 or 2876.1 EC: 1.11.1.9
6. PRDX6 was removed from the GPR’s associated with the following reactions: HMR_0960, HMR_0963, HMR_0988, HMR_2441, HMR_1048, HMR_1049. These are all glutathione peroxidase reactions, which PRDX6 is not involved in.
7. Reactions catalyzed by isoforms of peroxiredoxin were added: *Originally*: None *Updated*:
  ⍰ 1 h2o2[c] + 1 trdrd[c] → 2 h2o[c] + 1 trdox[c] GPR: 5052.1 or 5052.2 or 5052.3 or 7001.1 or 7001.2 or 7001.3 EC: 1.11.1.15
  ⍰ 1 h2o2[m] + 1 trdrd[m] → 2 h2o[m] + 1 trdox[m] GPR: 10935.1 or 10935.2 EC: 1.11.1.15
8. The oxidation and glutathionylation of protein thiol groups, as well as their reduction by glutaredoxin and thioredoxin, were added: *Originally*: None *Updated*:
  ⍰ 1 h2o2[c] + 1 Pr-SH[c] + 1 gthrd[c] → 2 h2o[c] + 1 Pr-SSG[c] GPR: - EC: -
  ⍰ 1 h2o2[m] + 1 Pr-SH[m] + 1 gthrd[m] → 2 h2o[m] + 1 Pr-SSG[m] GPR: - EC: -
  ⍰ 1 Pr-SSG[c] + 1 gthrd[c] → 1 Pr-SH[c] + 1 gthox[c] GPR: 2745.1 EC: 1.20.4.1
  ⍰ 1 Pr-SSG[m] + 1 gthrd[m] → 1 Pr-SH[m] + 1 gthox[m] GPR: 2745.1 EC: 1.20.4.1
  ⍰ 1 h2o2[c] + 1 Pr-SH[c] → 2 h2o[c] + 1 Pr-SS[c] GPR: - EC: -
  ⍰ 1 h2o2[m] + 1 Pr-SH[m] → 2 h2o[m] + 1 Pr-SS[m] GPR: - EC: -
  ⍰ 1 Pr-SS[c] + 1 trdrd[c] → 1 Pr-SH[c] + 1 trdox[c] GPR: - EC: -
  ⍰ 1 Pr-SS[m] + 1 trdrd[m] → 1 Pr-SH[m] + 1 trdox[m] GPR: - EC: -
9. The reliance of catalase enzyme on NADPH to prevent oxidation of its active site by H_2_O_2_ was incorporated (Kirkman et al., 1999). Kirkman et al. measured rates of H_2_O_2_ formation and corresponding rates of NADPH oxidation in mixtures of catalase and varying concentrations of glucose oxidase (Kirkman et al., 1987). If the rate of H_2_O_2_ formation by glucose oxidase is assumed to equal the rate of H_2_O_2_ clearance by catalase, then the slope of NADPH oxidation vs. H_2_O_2_ formation gives the amount of NADPH oxidized by catalase per molecule of H_2_O_2_ cleared. Linear regression of data from Kirkman et al. shows that for every 1 molecule of H_2_O_2_ cleared by catalase, 0.0641 molecules of NADPH are oxidized; thus, the chemical equation for catalase activity is:

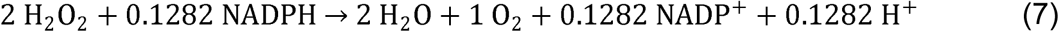 This change was made for catalase reactions in the cytosol, mitochondria, peroxisome, and endoplasmic reticulum.
10. The neomorphic 2-hydroxyglutarate-producing reaction catalyzed by mutant IDH1 was added. Reactions for 2-hydroxyglutarate export were also added. *Originally*: None *Updated*:
  ⍰ 1 akg[c] + 1 nadph[c] → 1 M00653[c] + 1 nadp[c] GPR: 3417.1 EC: 1.1.1.42
  - 1 M00653[c] → 1 M00653[e] GPR: None EC: None
  - 1 M00653[e] → GPR: None EC: None

Because transcriptomic and proteomic data from TCGA were available at the individual gene level but not at the individual isoform level, all isoforms for each individual gene within Recon3D were combined.

Other changes made to Recon3D include:

⍰ Removing the following duplicate reactions (when another reaction exists where the stoichiometric vector is an exact multiple of these duplicate reactions): “HMR_7257”, “PEHSFABPe”, “G6PDH2c”, “GNDc”, “PGLc”, “RPEc”
⍰ Removing EC numbers from all transport reactions, since many of these were incorrect.
⍰ Removing the following genes and any associations with them within Recon3D reactions, since information about these genes could not be found: NCBI Gene ID’s: 0, 100507855, 8041
⍰ Removing reactions within the following subsystems, as this was necessary to prevent de novo production of essential amino acids: “Protein assembly”, “Protein degradation”, “Protein modification”, “Protein formation”

- Removing reactions in the “Bile acid synthesis” subsystem, as these are not expected to be active in most tumors
- Removing NQO1 futile cycle reactions, as these un-constrained reactions resulted in erroneous predictions of NADPH production: “NADPQNOXR”, “NADQNOXR”, “HMR_9538”, “HMR_9722”
- Removing oxalosuccinic acid-producing IDH1 reactions, as these overlapped with α-ketoglutarate-producing IDH1 reactions: “r0423”, “r0424”, “r0425”, “r0422”

#### Model Flux Constraints

Upper flux bounds for reactions within Recon3D with both associated GPR rules and EC numbers were constrained using the Michaelis-Menten V_max_ parameter:

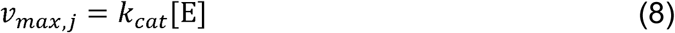

where *k*_*cat*_ and [E] are the turnover number (units of hr^-1^) and abundance (units of mmol gDW^-1^) of the associated enzyme catalyzing that reaction, respectively. Equation 8 was used to set upper flux bounds *v*_max,*j*_, as well as lower flux bounds *v*_min,*j*_ for reversible reactions (where *v*_min,*j*_ < 0). See sections “Enzyme Abundance Calculation” and “Turnover Number Calculation” for more information.

#### Enzyme Abundance Calculation

RNA-Seq gene expression data from TCGA patient tumors was obtained from Rahman et al.’s alternative preprocessing method (GEO: GSE62944) (Rahman et al., 2015). Data from this preprocessing method showed fewer missing values, more consistent expression between replicates, and improved prediction of biological pathway activity compared to the original TCGA pipeline.

Enzyme abundances within individual samples were predicted from sample gene expression data using the protein prediction pipeline shown in **Figure S1A**. The approach described below relies only on gene expression data from the individual sample of interest and not data from other samples included within the dataset, improving the reproducibility of this method as well as allowing for new samples to be analyzed post-hoc. The individual steps in the pipeline are explained herein:

1. *Schwanhäusser et al*. - By measuring the transcription, translation, and degradation rates of individual mRNA’s and proteins in NIH3T3 cells, Schwanhäusser et al. developed ODE models relating mRNA and protein abundances per cell that encompass 955/3268 (29.2%) genes in Recon3D (Schwanhausser et al., 2011). Their model also took into account proteins with half-lives longer than the length of the cell cycle, and how this would impact measurements of protein abundance. The authors demonstrated that these rate constants and models could be applied to accurately predict protein abundances in other samples including cancer cell lines. Using this model, gene expression values (transcripts per kilobase-million (TPM)-normalized) were converted to protein abundance values (parts per million (PPM)-normalized) for the 955 Recon3D genes included.
2. *Linear regression* - For each sample, a linear regression of predicted protein abundance values using the Schwanhäusser et al. method versus measured gene expression values of corresponding genes is performed (example is shown in **Figure S1B**). This regression model is used to predict the abundance of proteins where an associated gene expression value is available but not corresponding model/parameter values in the Schwanhäusser et al. method.
3. *PaxDB* - If an individual gene expression value is missing for a particular sample, the harmonic mean of abundance values for that associated protein across all *Homo sapiens* samples within the PaxDB database is taken (Wang et al., 2015). If the associated protein is not available in the PaxDB database, the harmonic mean of abundance values for all proteins across all *Homo sapiens* samples is taken. Imposing the same values for all samples with missing gene expression values acts to prevent artificial differences in FBA model predictions between samples due to missing data. The harmonic mean of PaxDB values is taken since (1) the arithmetic mean tends to overestimate representative values of positively-skewed distributions (as seen with protein abundance values); and (2) the smaller protein abundance estimate of the harmonic mean well characterizes the fact that missing RNA-Seq values are usually at least in part due to low actual mRNA expression.

For genes with associated enzymes in multiple cellular compartments (e.g. catalase, which localizes to the cytosol, mitochondria, peroxisome, and endoplasmic reticulum), the predicted protein expression value was divided into each cellular compartment with weights proportional to the COMPARTMENTS value for the cellular compartment in which the metabolic reaction takes place (Binder et al., 2014). These COMPARTMENTS values provide a score representing the confidence that a particular protein is found within a particular cellular compartment, using both experimental and computational evidence.

Validation of this pipeline was performed to ensure (1) that the predicted enzyme abundance values correlated better with experimental protein expression data compared to gene expression data; and (2) that the predicted enzyme abundance values were at physiological orders of magnitude. Normalized RPPA experimental protein expression values from TCGA samples were available for 30 genes/proteins in Recon3D (Weinstein et al., 2013). The correlation between experimental enzyme abundance and predicted enzyme abundance values (R^2^_PredictedAbundance_) was greater than the correlation between experimental enzyme abundance and gene expression values (R^2^_Gene_) for 18/30 genes (60%; **Figure S1C-E**). Additionally, the largest change in correlation for genes where R^2^_PredictedAbundance_ < R^2^_Gene_ was less than 0.004, indicating that the R^2^_PredictedAbundance_ was not substantially lower than the R^2^_Gene_ for any of the 30 genes.

Because experimental non-normalized protein abundance values were not available for TCGA samples for validation, experimentally-measured protein abundances in the NCI-60 panel of cancer cell lines were used to analyze the improvement in correlation to experimental data and physiological accuracy of predicted abundances on a genome scale. Experimental NCI-60 abundance values showed greater correlation to mean values of predicted enzyme abundances across TCGA samples (R^2^ = 0.450) compared to TCGA gene expression values (R^2^ = 0.314) (**Figure S1F-G**). Additionally, the order of magnitude of predicted abundance values matched well with experimental values for individual genes/proteins over the majority of orders of magnitude. The match to experimental values was greatest for enzyme abundances that were calculated at the first step of the pipeline (Schwanhäusser et al. method; R^2^ = 0.586) compared to abundances calculated at later steps (R^2^ = 0.286), indicating the necessity for future transcription, translation, and degradation rate measurements to be conducted for genes/proteins not currently covered in the Schwanhäusser et al. method (**Figure S1H-I**).

#### Turnover Number Calculation

Enzyme turnover numbers for associated reactions within Recon3D were determined using available experimental data from the BRENDA database API (Schomburg et al., 2004). Because the BRENDA database does not contain data for all Recon3D enzymes with the exact same enzyme commission (EC) number and substrate measured using human enzymes at 37°C and the correct cellular compartment-specific pH, a pipeline was developed where the most physiologically-accurate turnover numbers for Recon3D metabolic reactions are identified (**Figure S2**). For an individual enzyme-catalyzed reaction within Recon3D, the associated turnover number was determined as close to the start of the pipeline as data was available within BRENDA. Values determined towards the start of the pipeline are more accurate, but less turnover number data with this greater accuracy is available within BRENDA. In determining turnover number accuracy, priority was given in the order of (1) using the correct EC number (versus using data from EC numbers that match only the first 3, 2, or 1 digits with that of the Recon3D reaction); (2) using the correct substrate as given in the Recon3D reaction (versus using data from any available substrate); (3) using turnover number data taken from human enzymes (versus using data from any available organism); and (4) using turnover number data taken at physiological temperature and pH (versus using data near but not at physiological temperature and pH). At a particular step in the pipeline, if multiple turnover number values that met the pipeline criteria were available, a weighted average of these values was taken, with weights equal to the Tanimoto coefficient (a measure of molecular similarity; greater coefficient signifies greater similarity between two molecules) between the desired substrate in the Recon3D reaction and the substrate in BRENDA for which the associated turnover number was determined.

#### Mutation Data

The effect of single nucleotide polymorphisms (SNP’s) located in Recon3D genes on the catalytic activity of the associated metabolic enzyme was estimated using the Envision computational platform developed by Gray et al (Gray et al., 2018). Envision leverages large-scale experimental mutagenesis datasets to predict a score representing the effect of single amino acid changes on protein function; a score less than 1 signifies a loss-of-function, while a score greater than 1 signifies a gain-of-function. These quantitative scores were shown to correlate better with experimentally-measured percent changes in protein activity after mutation than previously-developed methods including SNAP2 and EVmutation. For each TCGA patient with mutation data available, the turnover number for a particular Recon3D reaction was multiplied by the Envision score for any SNP’s located in the associated gene.

#### IDH1 mutations

Particular mutations in isocitrate dehydrogenase 1 (IDH1), predominantly at amino acid R132, result in both loss of function of its normal NADPH-generating activity, as well as gain of function of a neomorphic NADPH-consuming activity that produces the oncometabolite 2-hydroxyglutarate (Dang et al., 2009). This change in function was implemented by adding the IDH1-catalyzed neomorphic reaction to Recon3D and imposing turnover numbers of the normal and neomorphic reactions measured by Avellaneda Matteo et al. based on the IDH1 mutation status of each tumor (**Table S3**) (Avellaneda Matteo et al., 2017).

#### Thermodynamic Constraints

Only metabolic reactions with a negative change in Gibbs free energy (ΔG) can carry a non-zero net flux. Implementing thermodynamic constraints prevents fluxes through these thermodynamically infeasible reactions; additionally, this limits the formation of thermodynamically infeasible loops that carry net fluxes around closed cycles in the metabolic network (Schellenberger et al., 2011). To implement thermodynamic constraints on FBA models, the minimum and maximum transformed Gibbs free energy of Recon3D reactions were obtained from the Virtual Metabolic Human database (Noor et al., 2013). For all reactions where both the minimum and maximum ΔG’^□^ values were greater than zero (and for all reversible reactions where both the minimum and maximum ΔG’^□^ values were less than zero in the reverse direction), the maximum flux *v*_max,*j*_ (or *v*_min,*j*_ for reversible reactions) through that reaction was set to zero.

#### Media Constraints

To simulate experimental cell culture conditions, the modeled external compartment contained all metabolites found in RPMI-1640 cell culture media as well as fetal bovine serum (Price and Gregory, 1982). The external compartment also contained water, hydrogen ions, hydroxide ions, oxygen, and carbon dioxide. Modeled samples were allowed to uptake these available metabolites but not any other metabolites found in Recon3D.

#### siRNA Transfections

Cells were seeded in 24-well plates at 5 × 10^4^ cells per well for glutathione half-cell potential measurements, or 96-well plates at 8 × 10^3^ cells per well for glucose oxidase response measurements. 24 hours after seeding, cells were transfected using the N-TER Nanoparticle siRNA Transfection System (Sigma-Aldrich; Cat#N2913) at a final siRNA concentration of 50 nM with serum-free medium for 4 hours; afterwards, an equal volume of 2x (20%) FBS-containing medium was added to each well. Negative controls consisted of transfection with the MISSION siRNA Universal Negative Control (Sigma-Aldrich; Cat#SIC001; Lot#WDAA1199). Predesigned MISSION siRNA’s targeting individual genes were ordered from Sigma-Aldrich; the top 3 rated siRNA’s based on expected efficacy for each gene target were pooled together and transfected concurrently, except for *GLUD1, GLUD2*, and *SUCLG2*, where only 2, 1, and 1 siRNA’s were available, respectively (**Table S4**). **Figure S5** shows the knockdown efficiency of siRNA transfections in each cell line with *GAPDH* siRNA compared to no siRNA (just N-TER transfection reagent) and negative control siRNA by Western blot (GAPDH antibody: Cell Signaling Technology, Cat#2118; Tubulin antibody: Thermo Fisher Scientific, Cat#62204). Experiments were performed 24hr after transfection.

#### E_hc_ GSH/GSSG Measurements

The protocol and reagents used for measurement of glutathione half-cell potential were adapted from Rahman et al. (Rahman et al., 2006). KPE buffer was made by combining 16mL of solution A (6.8g KH_2_PO_4_ (Sigma-Aldrich; Cat#P5655; CAS: 7778-77-0) in 500mL dH_2_O), 84mL of solution B (8.5g K_2_HPO_4_ (Sigma-Aldrich; Cat#P8281; CAS: 7758-11-4) in 500mL dH_2_O), and 0.327g EDTA sodium salt (Sigma-Aldrich; Cat#E5134; CAS: 6381-92-6). After removal of cell culture media, 150μL of extraction buffer (0.1% Trition-X100 (Sigma-Aldrich; Cat#X100; CAS: 9002-93-1) and 0.6% sulfosalicylic acid (Sigma-Aldrich; Cat#S2130; CAS: 5965-83-3) in KPE) was added to each well of a 24-well plate containing samples of interest. Plates were shaken at 800rpm for 10min to promote cell lysis. For each well, 100μL of lysate was taken for GSSG measurement, and 20μL was taken for GSSG+GSH measurement.

For GSSG+GSH measurement, 20μL samples were placed in a clear 96-well plate. 20μL of GSSG (Sigma-Aldrich; Cat#G6654; CAS: 27025-41-8) and GSH (Sigma-Aldrich; Cat#S4251; CAS: 70-18-8) standards at concentrations of 2e-2, 1e-2, 5e-3, 2.5e-3, 1e-3, 5e-4, 2.5e-4, 1e-4, 5e-5, 2.5e-5, 1e-5, and 0 mg/mL were placed in separate rows of the 96-well plate. 120μL of 1:1 DTNB:GR solution (DTNB: 2mg 5,5’-dithiobis(2-nitrobenzoic acid) (Sigma-Aldrich; Cat#D8130; CAS: 69-78-3) in 3mL KPE; GR: 40μL of glutathione reductase enzyme (Sigma-Aldrich; Cat#G3664; CAS: 9001-48-3) (250 units/mL) in 3mL KPE) was added to each well. 30sec later, 60μL of NADPH solution (2mg of β-NADPH (Sigma-Aldrich; Cat#N7505; CAS: 2646-71-1 (anhydrous)) in 3mL KPE) was added to each well. Immediately after, absorbance at 412nm in each well was measured every 31sec for a period of 5min10sec (11 measurements) using a BioTek Synergy 4 plate reader. All sample and standard values were background subtracted. To determine the concentration of GSSG+GSH in each sample (mg/mL), the slope of absorbance vs. time from each sample was compared to the average slope of the GSSG and GSH standard concentrations.

For GSSG measurement, 100μL samples as well as 100μL of GSSG and GSH standards were placed in a 96-well plate. 10μL of 2VP solution (2-vinylpyridine (Sigma-Aldrich; Cat#132292; CAS: 100-69-6) diluted 1:50 in KPE) was added to each well. 60min later, 10μL of triethanolamine solution (triethanolamine (Sigma-Aldrich; Cat#T58300; CAS: 102-71-6) diluted 1:10 in KPE) was added to each well. 10min later, 20μL from each well was transferred to a new clear 96-well plate. GSSG was then measured analogously to GSSG+GSH. GSSG measurements were multiplied by 1.2 to account for the dilution of the original 100μL sample with 10μL of 2VP solution and 10μL of triethanolamine solution.

To determine the concentration of GSH (mg/mL) for each sample, the measured concentration of GSSG was subtracted from the measured concentration of GSSG+GSH. To convert lysate concentrations (mg/mL) to intracellular concentrations (M), the following conversion was used:

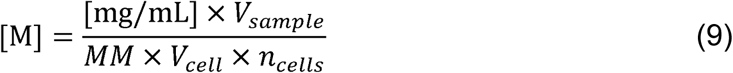

where *MM* is the molar mass of either GSH or GSSG (g/mol), *V*_*sample*_ is the volume of sample (20μL), *V*_*cell*_ is the estimated volume of each cell (1 pL), and *n*_*cells*_ is the number of cells per sample (5 × 10^4^).

The glutathione half-cell potential *E*_*hc*_ is calculated using the Nernst equation:

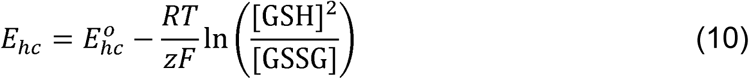

where *E*_*hc*_^*o*^ is the standard half-cell potential at pH 7.4 (−264 mV), *R* is the universal gas constant (8.314 J/K/mol), *T* is the temperature (310.15 K), *z* is the number of electrons (2), and *F* is the Faraday constant (96485 C/mol = 96.485 J/mV/mol).

#### Glucose Oxidase Response Measurements

Prior to seeding, cells were stained with CellTracker Red CMTPX Dye (Thermo Fisher Scientific; Cat#C34552) for 30min. 8 × 10^3^ cells per well were seeded in white-sided clear-bottom 96-well plates. 24hr after transfection and prior to treatment with glucose oxidase, CellTracker fluorescence measurements in each well were taken at Ex 577nm, Em 602nm using a BioTek Synergy 4 plate reader as a proxy for number of cells per well. An average of 10 fluorescence measurements was taken, and measurements were background subtracted. Samples were then treated with 10 mU/mL glucose oxidase enzyme (Sigma-Aldrich; Cat#G7141) diluted in 100μL cell culture medium (RPMI-1640 + 10% FBS) for 2hr. Afterwards, 100μL of CellTiter-Glo 2.0 Cell Viability Assay (Promega; Cat#G9241) was added to each well. Plates were shaken at 800rpm for 2min to promote cell lysis, and then incubated at room temperature for 10min. Luminescence measurements were then taken using a BioTek Synergy 4 plate reader and background subtracted. Cell viability normalized by cell count was measured by dividing the luminescence measurement by the average CellTracker fluorescence measurement.

### QUANTIFICATION AND STATISTICAL ANALYSIS

- *Comparison on objective values, reactions fluxes, and gene knockout effects between radiation-sensitive and -resistant FBA tumor models (Fig. 2A, 2D, 2F, 3B, 4A, 4C, S3, S4)*
  - n: number of individual FBA tumor models in each class. n = 716 radiation-sensitive tumor models, n = 199 radiation-resistant tumor models.
  - Boxplots: box = 25^th^, 50^th^, and 75^th^ percentiles, whiskers = 1.5 times the interquartile range.
  - Statistical test: 2-sided 2-sample Welch’s t-test without assumption of equal population variance.
- *Comparison of experimental glutathione half-cell potentials between radiation-sensitive and -resistant cell lines (Fig. 2C)*
  - n: number of biological replicates performed for each cell line. n = 12 for all cell lines except rSCC-61, where n = 10 (erroneously-negative GSSG values were obtained for 2/12 samples due to being below the limit of detection, and these were removed from the analysis).
  - Boxplots: box = 25^th^, 50^th^, and 75^th^ percentiles, whiskers = 1.5 times the interquartile range.
  - Statistical test: 2-sided 2-sample Welch’s t-test without assumption of equal population variance.
- *Comparison of gene knockout effects between radiation-sensitive and -resistant FBA tumor models (Fig. 3C, 3D, 4D)*
  - n: number of individual FBA tumor models in each class. n = 716 radiation-sensitive tumor models, n = 199 radiation-resistant tumor models.
  - Statistical test: 2-sided 2-sample Welch’s t-test without assumption of equal population variance. p-values for each of the 3,268 simulated gene knockouts performed were subsequently adjusted for multiple hypothesis testing using the Benjamini-Hochberg procedure for independent tests.
- *Comparison of model-predicted and experimentally-validated effect of gene knockdown on GSH production (Fig. 3F)*
  - n_1_: number of biological replicates performed for each cell line, for each gene knockdown and negative control. n_1_ = 3 for all cases
  - n_2_: number of cell line pairs for comparing to model predictions. n_2_ = 3
  - Statistical test: 2-sided 1-sample t-test with null hypothesis population mean = 0
- *Comparison of experimental glucose oxidase response between radiation-sensitive and -resistant cell lines (Fig. 4B)*
  - n: number of biological replicates performed for each cell line, for both 0 mU/mL and 10 mU/mL glucose oxidase treatment. Outliers were automatically detected and removed as determined by Grubbs’ 2-tailed test with α=0.05. n = 5 for GBM and HNSC cell lines, except for rSCC-61 10 mU/mL (n = 4) due to outlier removal. n = 10 for BRCA cell lines, except for NQO1(-) 0 mU/mL (n = 9) due to outlier removal.
  - Bar chart: mean ± 1 standard error.
  - Statistical test: 2-sided 2-sample t-test from mean, standard deviation, and number of observations from both samples
- *Comparison of model-predicted and experimentally-validated effect of gene knockdown on H*_*2*_*O*_*2*_ *response (Fig. 4F)*
  - n_1_: number of biological replicates performed for each cell line, for both glucose oxidase treatments, for each gene knockdown and negative control. Outliers were automatically detected and removed as determined by Grubbs’ 2-tailed test with α=0.05. n_1_ = 5 for GBM and HNSC cell lines, except for 1) M059K 10 mU/mL TKT KD (n=4) and 2) rSCC-61 10 mU/mL PGAM2 KD (n=4) due to outlier removal. n_1_ = 10 for BRCA cell lines, except for 1) NQO1(-) 0 mU/mL PGD KD (n=9) and 2) NQO1(-) 0 mU/mL TKT PD (n=9) due to outlier removal.
  - n_2_: number of cell line pairs for comparing to model predictions. n_2_ = 3
  - Statistical test: 2-sided 1-sample t-test with null hypothesis population mean = 0
- *Correlation between patient clinical factors and NADPH-generating fluxes in radiation-resistant tumor models (Fig. 5A)*
  - n: number of radiation-resistant tumor models. n = 199
  - Statistical test:
    ▪ Numerical factors: univariate regression
    ▪ Categorical factors: 1-way ANOVA

## REFERENCES

Adimora, N.J., Jones, D.P., and Kemp, M.L. (2010). A model of redox kinetics implicates the thiol proteome in cellular hydrogen peroxide responses. Antioxid Redox Signal 13, 731–743.

Agarwal, A.R., Yin, F., and Cadenas, E. (2014). Short-term cigarette smoke exposure leads to metabolic alterations in lung alveolar cells. Am J Respir Cell Mol Biol 51, 284–293.

Alvarez-Idaboy, J.R., and Galano, A. (2012). On the chemical repair of DNA radicals by glutathione: hydrogen vs electron transfer. J Phys Chem B 116, 9316–9325.

Avellaneda Matteo, D., Grunseth, A.J., Gonzalez, E.R., Anselmo, S.L., Kennedy, M.A., Moman, P., Scott, D.A., Hoang, A., and Sohl, C.D. (2017). Molecular mechanisms of isocitrate dehydrogenase 1 (IDH1) mutations identified in tumors: The role of size and hydrophobicity at residue 132 on catalytic efficiency. J Biol Chem 292, 7971–7983.

Bankar, S.B., Bule, M.V., Singhal, R.S., and Ananthanarayan, L. (2009). Glucose oxidase--an overview. Biotechnol Adv 27, 489–501.

Becker, S.A., and Palsson, B.O. (2008). Context-specific metabolic networks are consistent with experiments. PLoS Comput Biol 4, e1000082.

Benfeitas, R., Bidkhori, G., Mukhopadhyay, B., Klevstig, M., Arif, M., Zhang, C., Lee, S., Cinar, R., Nielsen, J., Uhlen, M., et al. (2019). Characterization of heterogeneous redox responses in hepatocellular carcinoma patients using network analysis. EBioMedicine 40, 471–487.

Binder, J.X., Pletscher-Frankild, S., Tsafou, K., Stolte, C., O’Donoghue, S.I., Schneider, R., and Jensen, L.J. (2014). COMPARTMENTS: unification and visualization of protein subcellular localization evidence. Database (Oxford) 2014, bau012.

Blazier, A.S., and Papin, J.A. (2012). Integration of expression data in genome-scale metabolic network reconstructions. Front Physiol 3, 299.

Bordel, S. (2018). Constraint based modeling of metabolism allows finding metabolic cancer hallmarks and identifying personalized therapeutic windows. Oncotarget 9, 19716–19729.

Brady, L.W., Perez, C.A., and Wazer, D.E. (2013). Perez & Brady’s principles and practice of radiation oncology. (Lippincott Williams & Wilkins).

Brunk, E., Sahoo, S., Zielinski, D.C., Altunkaya, A., Drager, A., Mih, N., Gatto, F., Nilsson, A., Preciat Gonzalez, G.A., Aurich, M.K., et al. (2018). Recon3D enables a three-dimensional view of gene variation in human metabolism. Nat Biotechnol 36, 272–281.

Cadet, J., and Wagner, J.R. (2013). DNA base damage by reactive oxygen species, oxidizing agents, and UV radiation. Cold Spring Harb Perspect Biol 5.

Cao, L., Li, L.S., Spruell, C., Xiao, L., Chakrabarti, G., Bey, E.A., Reinicke, K.E., Srougi, M.C., Moore, Z., and Dong, Y. (2014). Tumor-selective, futile redox cycle-induced bystander effects elicited by NQO1 bioactivatable radiosensitizing drugs in triple-negative breast cancers. Antioxidants & redox signaling 21, 237–250.

Chatterjee, A. (2013). Reduced glutathione: a radioprotector or a modulator of DNA-repair activity? Nutrients 5, 525–542.

Chen, X., Liu, L., Mims, J., Punska, E.C., Williams, K.E., Zhao, W., Arcaro, K.F., Tsang, A.W., Zhou, X., and Furdui, C.M. (2015). Analysis of DNA methylation and gene expression in radiation-resistant head and neck tumors. Epigenetics 10, 545–561.

Chen, X., Mims, J., Huang, X., Singh, N., Motea, E., Planchon, S.M., Beg, M., Tsang, A.W., Porosnicu, M., Kemp, M.L., et al. (2018). Modulators of Redox Metabolism in Head and Neck Cancer. Antioxid Redox Signal 29, 1660–1690.

Choi, Y.K., and Park, K.G. (2018). Targeting Glutamine Metabolism for Cancer Treatment. Biomol Ther (Seoul) 26, 19–28.

Choi, Y.M., Kim, H.K., Shim, W., Anwar, M.A., Kwon, J.W., Kwon, H.K., Kim, H.J., Jeong, H., Kim, H.M., Hwang, D., et al. (2015). Mechanism of Cisplatin-Induced Cytotoxicity Is Correlated to Impaired Metabolism Due to Mitochondrial ROS Generation. PLoS One 10, e0135083.

Ciccarese, F., and Ciminale, V. (2017). Escaping Death: Mitochondrial Redox Homeostasis in Cancer Cells. Front Oncol 7, 117.

Colijn, C., Brandes, A., Zucker, J., Lun, D.S., Weiner, B., Farhat, M.R., Cheng, T.Y., Moody, D.B., Murray, M., and Galagan, J.E. (2009). Interpreting expression data with metabolic flux models: predicting Mycobacterium tuberculosis mycolic acid production. PLoS Comput Biol 5, e1000489.

Covert, M.W., Xiao, N., Chen, T.J., and Karr, J.R. (2008). Integrating metabolic, transcriptional regulatory and signal transduction models in Escherichia coli. Bioinformatics 24, 2044–2050.

Dang, L., White, D.W., Gross, S., Bennett, B.D., Bittinger, M.A., Driggers, E.M., Fantin, V.R., Jang, H.G., Jin, S., Keenan, M.C., et al. (2009). Cancer-associated IDH1 mutations produce 2-hydroxyglutarate. Nature 462, 739–744.

Daniela, F.A., María, L.L.L., Bernard, I.A.S., Andrea, G.-S.A., Lizeth, L.E.C., and Luis, R.-P.J. (2015). Intracellular redox status and cell death induced by H2O2 in a human retinal epithelial cell line (Arpe-19). American Journal of BioScience 3, 93–113.

DeBerardinis, R.J., and Chandel, N.S. (2016). Fundamentals of cancer metabolism. Sci Adv 2, e1600200.

Delaney, G., Jacob, S., Featherstone, C., and Barton, M. (2005). The role of radiotherapy in cancer treatment: estimating optimal utilization from a review of evidence-based clinical guidelines. Cancer 104, 1129–1137.

Fan, J., Ye, J., Kamphorst, J.J., Shlomi, T., Thompson, C.B., and Rabinowitz, J.D. (2014). Quantitative flux analysis reveals folate-dependent NADPH production. Nature 510, 298–302.

Folger, O., Jerby, L., Frezza, C., Gottlieb, E., Ruppin, E., and Shlomi, T. (2011). Predicting selective drug targets in cancer through metabolic networks. Mol Syst Biol 7, 501.

Follia, L., Ferrero, G., Mandili, G., Beccuti, M., Giordano, D., Spadi, R., Satolli, M.A., Evangelista, A., Katayama, H., Hong, W., et al. (2019). Integrative Analysis of Novel Metabolic Subtypes in Pancreatic Cancer Fosters New Prognostic Biomarkers. Front Oncol 9, 115.

Forshaw, T.E., Holmila, R., Nelson, K.J., Lewis, J.E., Kemp, M.L., Tsang, A.W., Poole, L.B., Lowther, W.T., and Furdui, C.M. (2019). Peroxiredoxins in Cancer and Response to Radiation Therapies. Antioxidants (Basel) 8.

Franklin, D.A., He, Y., Leslie, P.L., Tikunov, A.P., Fenger, N., Macdonald, J.M., and Zhang, Y. (2016). p53 coordinates DNA repair with nucleotide synthesis by suppressing PFKFB3 expression and promoting the pentose phosphate pathway. Sci Rep 6, 38067.

Fritzemeier, C.J., Hartleb, D., Szappanos, B., Papp, B., and Lercher, M.J. (2017). Erroneous energy-generating cycles in published genome scale metabolic networks: Identification and removal. PLoS Comput Biol 13, e1005494.

Garcia Sanchez, C.E., and Torres Saez, R.G. (2014). Comparison and analysis of objective functions in flux balance analysis. Biotechnol Prog 30, 985–991.

Gray, V.E., Hause, R.J., Luebeck, J., Shendure, J., and Fowler, D.M. (2018). Quantitative Missense Variant Effect Prediction Using Large-Scale Mutagenesis Data. Cell Syst 6, 116-124.e113.

Gujar, A.D., Le, S., Mao, D.D., Dadey, D.Y., Turski, A., Sasaki, Y., Aum, D., Luo, J., Dahiya, S., Yuan, L., et al. (2016). An NAD+-dependent transcriptional program governs self-renewal and radiation resistance in glioblastoma. Proc Natl Acad Sci U S A 113, E8247–e8256.

Harris, I.S., Treloar, A.E., Inoue, S., Sasaki, M., Gorrini, C., Lee, K.C., Yung, K.Y., Brenner, D., Knobbe-Thomsen, C.B., Cox, M.A., et al. (2015). Glutathione and thioredoxin antioxidant pathways synergize to drive cancer initiation and progression. Cancer Cell 27, 211–222.

Henry, C.S., Broadbelt, L.J., and Hatzimanikatis, V. (2007). Thermodynamics-based metabolic flux analysis. Biophys J 92, 1792–1805.

Hsieh, J.Y., Li, S.Y., Tsai, W.C., Liu, J.H., Lin, C.L., Liu, G.Y., and Hung, H.C. (2015). A small-molecule inhibitor suppresses the tumor-associated mitochondrial NAD(P)+-dependent malic enzyme (ME2) and induces cellular senescence. Oncotarget 6, 20084–20098.

Jaramillo, M.C., and Zhang, D.D. (2013). The emerging role of the Nrf2-Keap1 signaling pathway in cancer. Genes Dev 27, 2179–2191.

Jensen, P.A., and Papin, J.A. (2011). Functional integration of a metabolic network model and expression data without arbitrary thresholding. Bioinformatics 27, 541–547.

Jin, L., Li, D., Alesi, G.N., Fan, J., Kang, H.B., Lu, Z., Boggon, T.J., Jin, P., Yi, H., Wright, E.R., et al. (2015). Glutamate dehydrogenase 1 signals through antioxidant glutathione peroxidase 1 to regulate redox homeostasis and tumor growth. Cancer Cell 27, 257–270.

Jones, M.E. (1980). Pyrimidine nucleotide biosynthesis in animals: genes, enzymes, and regulation of UMP biosynthesis. Annu Rev Biochem 49, 253–279.

Kanarek, N., Keys, H.R., Cantor, J.R., Lewis, C.A., Chan, S.H., Kunchok, T., Abu-Remaileh, M., Freinkman, E., Schweitzer, L.D., and Sabatini, D.M. (2018). Histidine catabolism is a major determinant of methotrexate sensitivity. Nature 559, 632–636.

Kanarek, N., Petrova, B., and Sabatini, D.M. (2020). Dietary modifications for enhanced cancer therapy. Nature 579, 507–517.

Kansanen, E., Kuosmanen, S.M., Leinonen, H., and Levonen, A.L. (2013). The Keap1-Nrf2 pathway: Mechanisms of activation and dysregulation in cancer. Redox Biol 1, 45–49.

Kaushik, A.K., and DeBerardinis, R.J. (2018). Applications of metabolomics to study cancer metabolism. Biochim Biophys Acta Rev Cancer 1870, 2–14.

Kilburn, D.G., Lilly, M.D., and Webb, F.C. (1969). The energetics of mammalian cell growth. J Cell Sci 4, 645–654.

Kim, J., and DeBerardinis, R.J. (2019). Mechanisms and Implications of Metabolic Heterogeneity in Cancer. Cell Metab 30, 434–446.

Kirkman, H.N., Galiano, S., and Gaetani, G.F. (1987). The function of catalase-bound NADPH. J Biol Chem 262, 660–666.

Kirkman, H.N., Rolfo, M., Ferraris, A.M., and Gaetani, G.F. (1999). Mechanisms of protection of catalase by NADPH. Kinetics and stoichiometry. J Biol Chem 274, 13908–13914.

Lee, J.M., Gianchandani, E.P., Eddy, J.A., and Papin, J.A. (2008). Dynamic analysis of integrated signaling, metabolic, and regulatory networks. PLoS Comput Biol 4, e1000086.

Lee, Y.S., Oh, J.H., Yoon, S., Kwon, M.S., Song, C.W., Kim, K.H., Cho, M.J., Mollah, M.L., Je, Y.J., Kim, Y.D., et al. (2010). Differential gene expression profiles of radioresistant non-small-cell lung cancer cell lines established by fractionated irradiation: tumor protein p53-inducible protein 3 confers sensitivity to ionizing radiation. Int J Radiat Oncol Biol Phys 77, 858–866.

Lewis, C.A., Parker, S.J., Fiske, B.P., McCloskey, D., Gui, D.Y., Green, C.R., Vokes, N.I., Feist, A.M., Vander Heiden, M.G., and Metallo, C.M. (2014). Tracing compartmentalized NADPH metabolism in the cytosol and mitochondria of mammalian cells. Mol Cell 55, 253–263.

Lewis, J.E., Costantini, F., Mims, J., Chen, X., Furdui, C.M., Boothman, D.A., and Kemp, M.L. (2018). Genome-Scale Modeling of NADPH-Driven beta-Lapachone Sensitization in Head and Neck Squamous Cell Carcinoma. Antioxid Redox Signal 29, 937–952.

Lewis, J.E., Singh, N., Holmila, R.J., Sumer, B.D., Williams, N.S., Furdui, C.M., Kemp, M.L., and Boothman, D.A. (2019). Targeting NAD(+) Metabolism to Enhance Radiation Therapy Responses. Semin Radiat Oncol 29, 6–15.

Lin, T.-Y., Cantley, L.C., and DeNicola, G.M. (2016). NRF2 rewires cellular metabolism to support the antioxidant response. A Master Regulator of Oxidative Stress-The Transcription Factor Nrf2.

Mallikarjun, V., Clarke, D.J., and Campbell, C.J. (2012). Cellular redox potential and the biomolecular electrochemical series: a systems hypothesis. Free Radical Biology and Medicine 53, 280–288.

Manem, V.S., and Dhawan, A. (2019). RadiationGeneSigDB: a database of oxic and hypoxic radiation response gene signatures and their utility in pre-clinical research. Br J Radiol 92, 20190198.

Marullo, R., Werner, E., Degtyareva, N., Moore, B., Altavilla, G., Ramalingam, S.S., and Doetsch, P.W. (2013). Cisplatin induces a mitochondrial-ROS response that contributes to cytotoxicity depending on mitochondrial redox status and bioenergetic functions. PLoS One 8, e81162.

Miller, K.D., Siegel, R.L., Lin, C.C., Mariotto, A.B., Kramer, J.L., Rowland, J.H., Stein, K.D., Alteri, R., and Jemal, A. (2016). Cancer treatment and survivorship statistics, 2016. CA Cancer J Clin 66, 271–289.

Mims, J., Bansal, N., Bharadwaj, M.S., Chen, X., Molina, A.J., Tsang, A.W., and Furdui, C.M. (2015). Energy metabolism in a matched model of radiation resistance for head and neck squamous cell cancer. Radiat Res 183, 291–304.

Nilsson, A., and Nielsen, J. (2017). Genome scale metabolic modeling of cancer. Metab Eng 43, 103–112.

Noor, E., Haraldsdottir, H.S., Milo, R., and Fleming, R.M. (2013). Consistent estimation of Gibbs energy using component contributions. PLoS Comput Biol 9, e1003098.

Noronha-Dutra, A.A., Epperlein, M.M., and Woolf, N. (1993). Effect of cigarette smoking on cultured human endothelial cells. Cardiovasc Res 27, 774–778.

Oberhardt, M.A., Goldberg, J.B., Hogardt, M., and Papin, J.A. (2010). Metabolic network analysis of Pseudomonas aeruginosa during chronic cystic fibrosis lung infection. Journal of bacteriology 192, 5534–5548.

Orth, J.D., Thiele, I., and Palsson, B.O. (2010). What is flux balance analysis? Nat Biotechnol 28, 245–248.

Porporato, P.E., Filigheddu, N., Pedro, J.M.B., Kroemer, G., and Galluzzi, L. (2018). Mitochondrial metabolism and cancer. Cell Res 28, 265–280.

Price, P.J., and Gregory, E.A. (1982). Relationship between in vitro growth promotion and biophysical and biochemical properties of the serum supplement. In Vitro 18, 576–584.

Rahman, I., Kode, A., and Biswas, S.K. (2006). Assay for quantitative determination of glutathione and glutathione disulfide levels using enzymatic recycling method. Nat Protoc 1, 3159–3165.

Rahman, M., Jackson, L.K., Johnson, W.E., Li, D.Y., Bild, A.H., and Piccolo, S.R. (2015). Alternative preprocessing of RNA-Sequencing data in The Cancer Genome Atlas leads to improved analysis results. Bioinformatics 31, 3666–3672.

Reisz, J.A., Bansal, N., Qian, J., Zhao, W., and Furdui, C.M. (2014). Effects of ionizing radiation on biological molecules--mechanisms of damage and emerging methods of detection. Antioxid Redox Signal 21, 260–292.

Schellenberger, J., Lewis, N.E., and Palsson, B.O. (2011). Elimination of thermodynamically infeasible loops in steady-state metabolic models. Biophys J 100, 544–553.

Schnell, J.R., Dyson, H.J., and Wright, P.E. (2004). Structure, dynamics, and catalytic function of dihydrofolate reductase. Annu Rev Biophys Biomol Struct 33, 119–140.

Schomburg, I., Chang, A., Ebeling, C., Gremse, M., Heldt, C., Huhn, G., and Schomburg, D. (2004). BRENDA, the enzyme database: updates and major new developments. Nucleic Acids Res 32, D431–433.

Schultz, A., and Qutub, A.A. (2016). Reconstruction of Tissue-Specific Metabolic Networks Using CORDA. PLoS Comput Biol 12, e1004808.

Schwanhausser, B., Busse, D., Li, N., Dittmar, G., Schuchhardt, J., Wolf, J., Chen, W., and Selbach, M. (2011). Global quantification of mammalian gene expression control. Nature 473, 337–342.

Shlomi, T., Cabili, M.N., Herrgard, M.J., Palsson, B.O., and Ruppin, E. (2008). Network-based prediction of human tissue-specific metabolism. Nat Biotechnol 26, 1003–1010.

Skvortsov, S., Debbage, P., Cho, W.C., Lukas, P., and Skvortsova, I. (2014). Putative biomarkers and therapeutic targets associated with radiation resistance. Expert Rev Proteomics 11, 207–214.

Skvortsova, I., Skvortsov, S., Stasyk, T., Raju, U., Popper, B.A., Schiestl, B., von Guggenberg, E., Neher, A., Bonn, G.K., Huber, L.A., et al. (2008). Intracellular signaling pathways regulating radioresistance of human prostate carcinoma cells. Proteomics 8, 4521–4533.

Smith, L., Qutob, O., Watson, M.B., Beavis, A.W., Potts, D., Welham, K.J., Garimella, V., Lind, M.J., Drew, P.J., and Cawkwell, L. (2009). Proteomic identification of putative biomarkers of radiotherapy resistance: a possible role for the 26S proteasome? Neoplasia 11, 1194–1207.

Spitz, D.R., Azzam, E.I., Li, J.J., and Gius, D. (2004). Metabolic oxidation/reduction reactions and cellular responses to ionizing radiation: a unifying concept in stress response biology. Cancer Metastasis Rev 23, 311–322.

Stein, E.M., Altman, J.K., Collins, R., DeAngelo, D.J., Fathi, A.T., Flinn, I., Frankel, A., Levine, R.L., Medeiros, B.C., and Patel, M. (2014). AG-221, an oral, selective, first-in-class, potent inhibitor of the IDH2 mutant metabolic enzyme, induces durable remissions in a phase I study in patients with IDH2 mutation positive advanced hematologic malignancies. (American Society of Hematology Washington, DC).

Stuani, L., Sabatier, M., and Sarry, J.E. (2019). Exploiting metabolic vulnerabilities for personalized therapy in acute myeloid leukemia. BMC Biol 17, 57.

Supandi, F., and Van Beek, J.H. (2018). Computational prediction of changes in brain metabolic fluxes during Parkinson’s disease from mRNA expression. PloS one 13.

Tominaga, H., Kodama, S., Matsuda, N., Suzuki, K., and Watanabe, M. (2004). Involvement of reactive oxygen species (ROS) in the induction of genetic instability by radiation. J Radiat Res 45, 181–188.

Turgeon, M.O., Perry, N.J.S., and Poulogiannis, G. (2018). DNA Damage, Repair, and Cancer Metabolism. Front Oncol 8, 15.

Wallace, T.C., Bultman, S., D’Adamo, C., Daniel, C.R., Debelius, J., Ho, E., Eliassen, H., Lemanne, D., Mukherjee, P., Seyfried, T.N., et al. (2019). Personalized Nutrition in Disrupting Cancer - Proceedings From the 2017 American College of Nutrition Annual Meeting. J Am Coll Nutr 38, 1–14.

Wang, M., Herrmann, C.J., Simonovic, M., Szklarczyk, D., and von Mering, C. (2015). Version 4.0 of PaxDb: Protein abundance data, integrated across model organisms, tissues, and cell-lines. Proteomics 15, 3163–3168.

Weinstein, J.N., Collisson, E.A., Mills, G.B., Shaw, K.R., Ozenberger, B.A., Ellrott, K., Shmulevich, I., Sander, C., and Stuart, J.M. (2013). The Cancer Genome Atlas Pan-Cancer analysis project. Nat Genet 45, 1113–1120.

Xiao, W., Wang, R.S., Handy, D.E., and Loscalzo, J. (2018). NAD(H) and NADP(H) Redox Couples and Cellular Energy Metabolism. Antioxid Redox Signal 28, 251–272.

Yin, F., Sancheti, H., and Cadenas, E. (2012). Silencing of nicotinamide nucleotide transhydrogenase impairs cellular redox homeostasis and energy metabolism in PC12 cells. Biochim Biophys Acta 1817, 401–409.

Yizhak, K., Le Dévédec, S.E., Rogkoti, V.M., Baenke, F., de Boer, V.C., Frezza, C., Schulze, A., van de Water, B., and Ruppin, E. (2014). A computational study of the Warburg effect identifies metabolic targets inhibiting cancer migration. Molecular systems biology 10.

Zhang, C., and Hua, Q. (2015). Applications of Genome-Scale Metabolic Models in Biotechnology and Systems Medicine. Front Physiol 6, 413.

Zhuang, H., Li, Q., Zhang, X., Ma, X., Wang, Z., Liu, Y., Yi, X., Chen, R., Han, F., Zhang, N., et al. (2018). Downregulation of glycine decarboxylase enhanced cofilin-mediated migration in hepatocellular carcinoma cells. Free Radic Biol Med 120, 1–12.

