## Supplemental Information for "Personalized Genome-Scale Metabolic Models Identify Targets of Redox Metabolism in Radiation-Resistant Tumors"

### SUPPLEMENTAL FIGURES

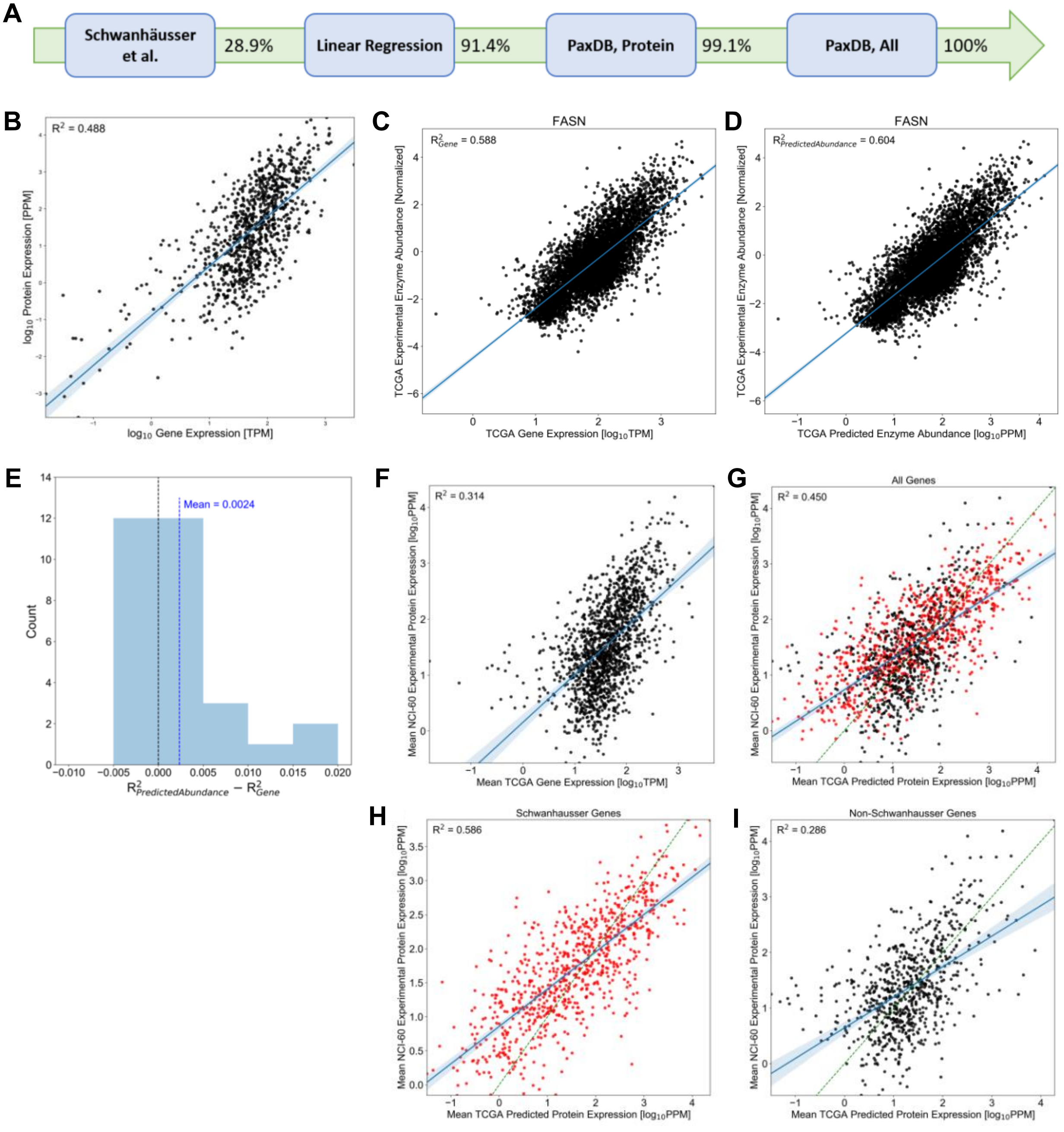


**Figure S1.** *Prediction of enzyme abundance values for FBA models of TCGA patient tumors (Related to **Figure 1**, **Methods**).* (**A**) Pipeline for the estimation of enzyme abundance values from gene expression data for individual TCGA patient tumors.

Percentages represent the percentage of enzyme abundances of associated genes in Recon3D that have been cumulatively determined at each step in the pipeline for all TCGA samples. (**B**) Example of linear regression step in the pipeline for enzyme abundance values, where predicted protein abundances using the Schwanhäusser et al. method are regressed versus the measured gene expression values of corresponding genes in an individual TCGA sample. (**C-D**) Example comparison of (C) gene expression and (D) predicted enzyme abundance values to experimental enzyme abundances from TCGA samples for an individual gene/protein (FASN). Blue line: least squares linear regression line. (**E**) Improvement of correlation to experimental enzyme abundances of predicted abundance values ($R^2_{PredictedAbundance}$) compared to original gene expression data ($R^2_{gene}$) for all 30 genes/proteins with available experimental values for TCGA samples. Blue dashed line: mean value of $R^2_{PredictedAbundance}$-$R^2_{gene}$ across all 30 genes/proteins. (**F**) Correlation between mean gene expression values across TCGA samples and mean experimental enzyme abundances across NCI-60 samples for all 3,268 genes/proteins in Recon3D. Blue line: least squares linear regression line. (**G**) Correlation between mean predicted enzyme abundances across TCGA samples and mean experimental enzyme abundances across NCI-60 samples for all 3,268 genes/proteins in Recon3D. This plot is broken down into (**H**; red dots) those values calculated during the first step of the prediction pipeline (Schwanhäusser et al. method) and (**I**; black dots) those values calculated at later steps of the pipeline. Green dashed line: 1:1 line.

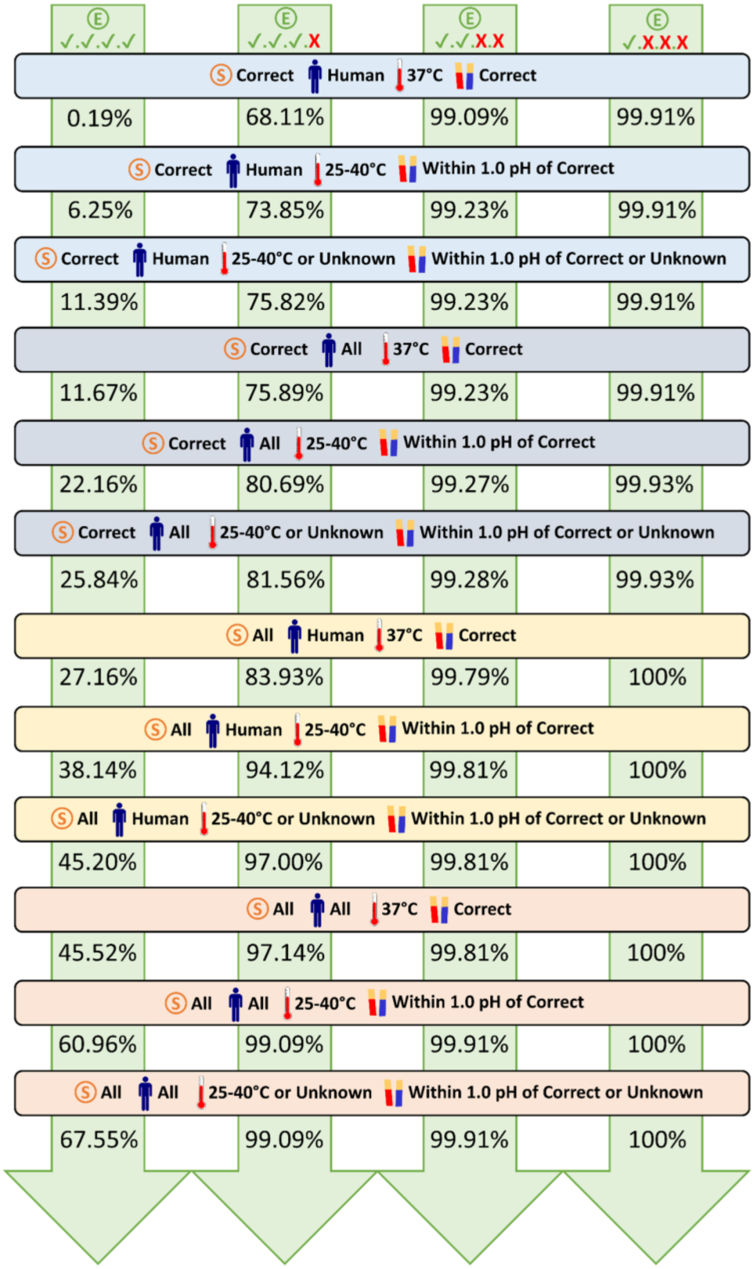


**Figure S2.** *Pipeline for the estimation of enzyme turnover numbers using data from the BRENDA database (Related **to Figure 1**, **Methods**).* Percentages represent the percentage of turnover numbers of associated reactions in Recon3D that have been cumulatively determined at each step in the pipeline. (E): Enzyme commission (EC) number. (S): Substrate.

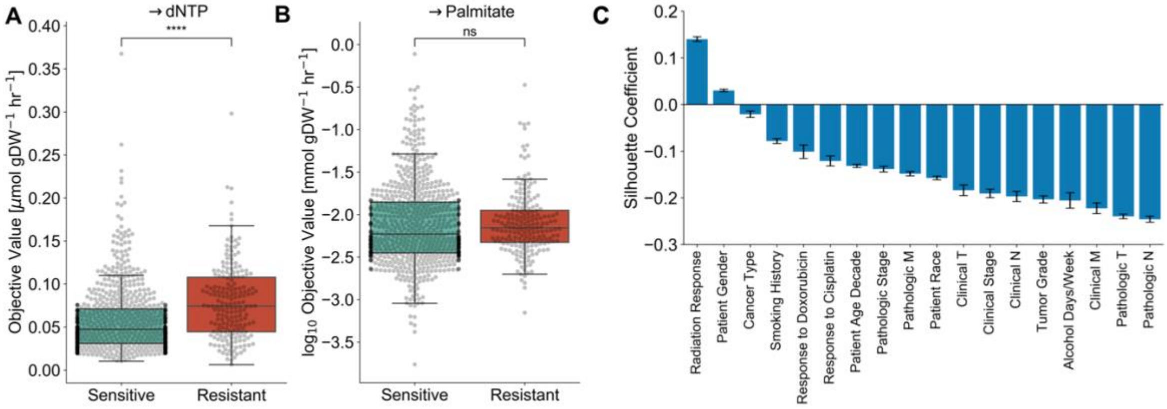


**Figure S3.** *Downstream effects of FBA-predicted NADPH production and correlation with patient clinical factors (Related to **Figure 2**).* (**A**) Comparison of FBA-predicted production of deoxynucleotides between radiation-sensitive and -resistant TCGA tumors. (**B**) Comparison of FBA-predicted production of palmitate between radiation-sensitive and -resistant TCGA tumors. (**C**) Silhouette coefficient between samples comparing radiation response and other clinical factors as cluster labels. For each factor, a more positive average silhouette coefficient signifies that samples cluster strongly together based on their factor value, and a more negative average silhouette coefficient signifies that samples cluster weakly based on their factor value (and that some other factor is likely to be the true separating factor). ns: not significant, *: $p \leq 0.05$, **: $p \leq 0.01$, ***: $p \leq 0.001$, ****: $p \leq 0.0001$.

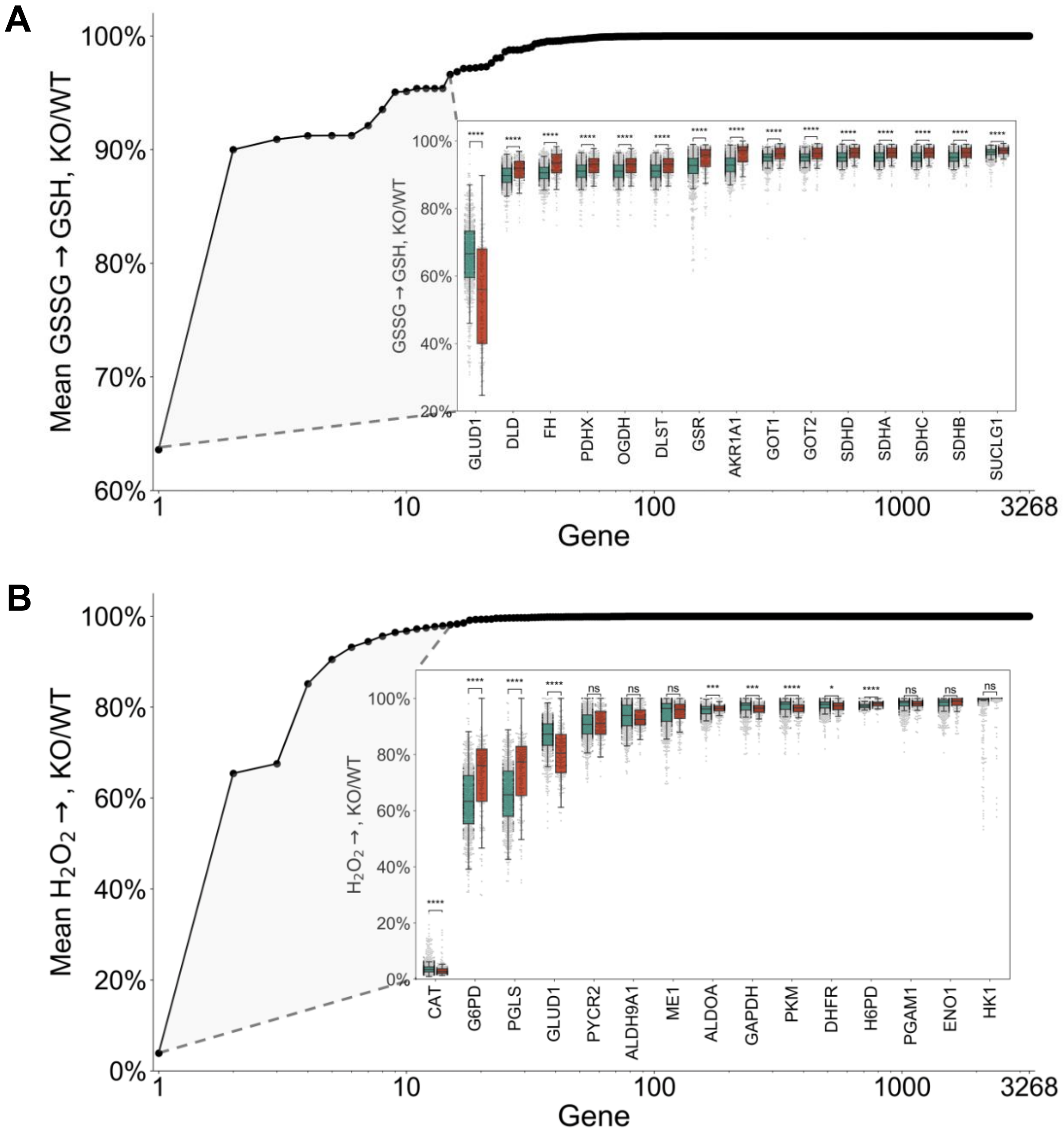


**Figure S4.** *Simulated genome-wide knockout screen for targets impacting GSH production and $H_2O_2$ clearance (Related to* ***Figures 3-4****).* (**A**) Effect of simulated knockout of each individual gene in Recon3D on total GSH production in TCGA tumors. Values are the ratio of total GSH production after versus before knockout. Genes are rank ordered based on increasing mean KO/WT ratio (decreasing gene knockout effect) across all tumor models. Outset: KO/WT ratios are averaged across all tumor models. Inset: For the top 15 genes, KO/WT ratios from individual patient tumor models are shown, along with the comparison between radiation-sensitive and -resistant cohorts. (**B**) Effect of simulated knockout of each individual gene in Recon3D on total $H_2O_2$ clearance in TCGA tumors. Values are the ratio of total $H_2O_2$ clearance after versus before knockout. Genes are rank ordered based on increasing mean KO/WT ratio (decreasing gene knockout effect) across all tumor models. Outset: KO/WT ratios are averaged across all tumor models. Inset: For the top 15 genes, KO/WT ratios from individual patient tumor models are shown, along with the comparison between radiation-sensitive and -resistant cohorts. ns: not significant, *: $p \leq 0.05$, **: $p \leq 0.01$, ***: $p \leq 0.001$, ****: $p \leq 0.0001$.

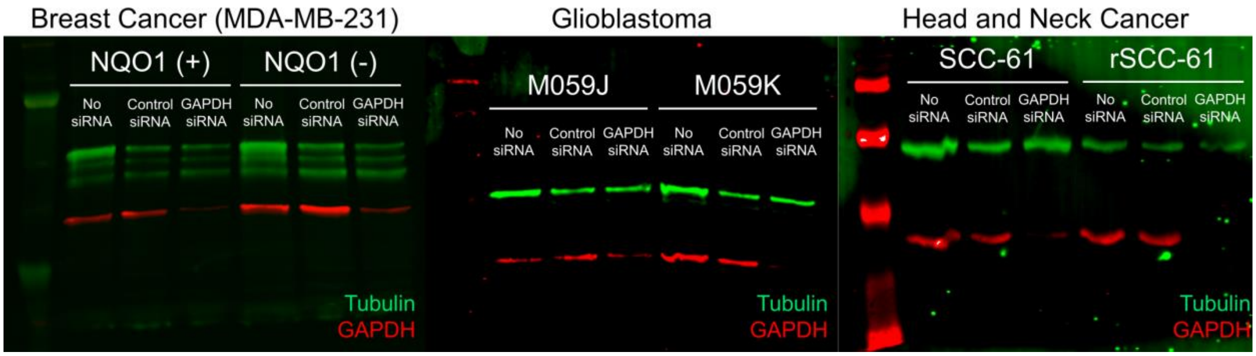


**Figure S5.** *Knockdown efficiency of siRNA transfections in each cell line with GAPDH (red) siRNA compared to no siRNA (just N-TER transfection reagent) and negative control siRNA (Related to* ***Methods****).* Tubulin (green) was used as a loading control.

### SUPPLEMENTAL TABLES

**Table S1.** *Matched radiation-sensitive and radiation-resistant cell lines (Related to* ***Figure 2****,* ***Methods****).*

| Cancer Type | Radiation-Sensitive | Radiation-Resistant | Notes | Source |
|---|---|---|---|---|
| Breast (BRCA) | MDA-MB-231 NQO1(-) (Sex: F) | MDA-MB-231 NQO1(+) (Sex: F) | Stable NQO1 expression was restored in NQO1(-) cells to create NQO1(+) cells. | Dr. David Boothman, Indiana University (Huang et al., 2016) |
| Glioblastoma (GBM) | M059J (Sex: M) | M059K (Sex: M) | Both isolated from same tumor specimen. M059J cells lack DNA-PK activity, rendering them more radiation-sensitive. | ATCC |
| Head and Neck (HNSC) | SCC-61 (Sex: M) | rSCC-61 (Sex: M) | rSCC-61 cells were derived from SCC-61 cells after repeated radiation exposure and selection of surviving colonies. | Dr. Cristina Furdui, Wake Forest University (Bansal et al., 2014; Zhao et al., 2011) |

**Table S2.** *Objective functions used in FBA and FVA (Related to* ***Methods****).*

| Title | Objective Function |
|---|---|
| $NADP^+$ → NADPH | 1 nadph[all] → 1 nadp[all] + 1 h[all] |
| $NADP^+$ → NADPH, Cytosolic | 1 nadph[c] → 1 nadp[c] + 1 h[c] |
| $NADP^+$ → NADPH, Mitochondrial | 1 nadph[m] → 1 nadp[m] + 1 h[m] |
| $NAD^+$ → NADH | 1 nadh[all] → 1 nad[all] + 1 h[all] |
| GSSG → GSH | 2 gthrd[all] + 1 nadp[all] → 1 gthox[all] + 1 nadph[all] + 1 h[all] |
| $H_2O_2$ → | Ø → 1 h2o2[all] |
| → dNTP | 1 datp[all] + 1 dctp[all] + 1 dgtp[all] + 1 dttp[all] → Ø |
| → Palmitate | 1 hdca[all] → Ø |

*Note*: “[all]” signifies that the objective function was maximized in all cellular compartments with equal weights. “Ø” signifies no metabolites on the left or right side of the equation.

**Table S3.** *Turnover numbers of normal and neomorphic IDH1 reactions for given IDH1 mutations (Related to* ***Methods****).*

| **Mutation** | **$k_{cat}$ Normal [$s^{-1}$]** | **$k_{cat}$ Neomorphic [$s^{-1}$]** |
|---|---|---|
| WT | 85 ± 4 | 0.019 ± 0.001 |
| R132C | 4.4 ± 0.1 | 1.60 ± 0.07 |
| R132G | 9.3 ± 0.6 | 1.59 ± 0.09 |
| R132H | 2.4 ± 0.1 | 4.2 ± 0.3 |

**Table S4.** *siRNA's used for each gene target (Related to* ***Methods****).*

| Gene Target | siRNA 1 | siRNA 2 | siRNA 3 |
|---|---|---|---|
| *ALDH1L1* | SASI_Hs01_00106766 | SASI_Hs01_00106767 | SASI_Hs01_00106768 |
| *ALDH4A1* | SASI_Hs01_00242114 | SASI_Hs01_00242115 | SASI_Hs01_00242116 |
| *ALDOA* | SASI_Hs01_00211472 | SASI_Hs01_00211474 | SASI_Hs01_00211476 |
| *CAT* | SASI_Hs02_00332471 | SASI_Hs01_00092507 | SASI_Hs01_00092508 |
| *CPT2* | SASI_Hs01_00121386 | SASI_Hs01_00121387 | SASI_Hs01_00121388 |
| *DLST* | SASI_Hs01_00176880 | SASI_Hs01_00176881 | SASI_Hs01_00176882 |
| *FH* | SASI_Hs01_00037389 | SASI_Hs01_00037390 | SASI_Hs01_00037391 |
| *G6PD* | SASI_Hs01_00013421 | SASI_Hs01_00013422 | SASI_Hs02_00302480 |
| *GLUD1* | SASI_Hs01_00082304 | SASI_Hs01_00082305 | |
| *GLUD2* | SASI_Hs01_00044099 | | |
| *GSR* | SASI_Hs01_00152913 | SASI_Hs01_00152914 | SASI_Hs02_00302987 |
| *LDHB* | SASI_Hs01_00181812 | SASI_Hs01_00181813 | SASI_Hs02_00333601 |
| *MTHFD1* | SASI_Hs01_00090500 | SASI_Hs01_00090501 | SASI_Hs01_00090502 |
| *MTHFR* | SASI_Hs01_00228224 | SASI_Hs01_00228225 | SASI_Hs01_00228226 |
| *NMNAT3* | SASI_Hs01_00068505 | SASI_Hs01_00068508 | SASI_Hs01_00068509 |
| *OGDH* | SASI_Hs01_00171350 | SASI_Hs01_00171351 | SASI_Hs01_00171353 |
| *PDHB* | SASI_Hs01_00164768 | SASI_Hs01_00164769 | SASI_Hs01_00164770 |
| *PGAM2* | SASI_Hs01_00037889 | SASI_Hs02_00302252 | SASI_Hs01_00037890 |
| *PGD* | SASI_Hs01_00134091 | SASI_Hs02_00334150 | SASI_Hs01_00134092 |
| *PRDX6* | SASI_Hs02_00338555 | SASI_Hs01_00182363 | SASI_Hs01_00182364 |
| *SHDA* | SASI_Hs02_00337042 | SASI_Hs01_00131081 | SASI_Hs02_00337043 |
| *SUCLG2* | SASI_Hs01_00106317 | | |
| *TKT* | SASI_Hs01_00241972 | SASI_Hs01_00241973 | SASI_Hs01_00241971 |
| *TPI1* | SASI_Hs01_00014384 | SASI_Hs01_00014385 | SASI_Hs01_00014386 |

**References:**


Bansal, N., Mims, J., Kuremsky, J.G., Olex, A.L., Zhao, W., Yin, L., Wani, R., Qian, J., Center, B., Marrs, G.S., et al. (2014). Broad phenotypic changes associated with gain of radiation resistance in head and neck squamous cell cancer. Antioxid Redox Signal *21*, 221-236.

Huang, X., Motea, E.A., Moore, Z.R., Yao, J., Dong, Y., Chakrabarti, G., Kilgore, J.A., Silvers, M.A., Patidar, P.L., Cholka, A., et al. (2016). Leveraging an NQO1 Bioactivatable Drug for Tumor-Selective Use of Poly(ADP-ribose) Polymerase Inhibitors. Cancer Cell *30*, 940-952.

Zhao, M., Sano, D., Pickering, C.R., Jasser, S.A., Henderson, Y.C., Clayman, G.L., Sturgis, E.M., Ow, T.J., Lotan, R., Carey, T.E., et al. (2011). Assembly and initial characterization of a panel of 85 genomically validated cell lines from diverse head and neck tumor sites. Clin Cancer Res *17*, 7248-7264.
